# Conserved Role of the Large Conductance Calcium-Activated Potassium Channel, K_Ca_1.1, in Sinus Node Function and Arrhythmia Risk

**DOI:** 10.1101/2020.06.28.176495

**Authors:** Santiago Pineda, Vesna Nikolova-Krstevski, Christiana Leimena, Andrew J. Atkinson, Ann-Kristin Altekoester, Charles D. Cox, Arie Jacoby, Inken G. Huttner, Yue-Kun Ju, Magdalena Soka, Monique Ohanian, Gunjan Trivedi, Sreehari Kalvakuri, Renee Johnson, Peter Molenaar, Dennis Kuchar, David G. Allen, Dirk F. van Helden, Richard P. Harvey, Adam P. Hill, Rolf Bodmer, Georg Vogler, Halina Dobrzynski, Karen Ocorr, Diane Fatkin

**Author notes:** Correspondence: Diane Fatkin, MD, Victor Chang Cardiac Research Institute, Lowy Packer Building, 405 Liverpool St, PO Box 699, Darlinghurst NSW 2010, Australia; or Karen Ocorr, PhD, Sanford-Burnham Prebys Medical Discovery Institute, 10901 N. Torrey Pines Road, La Jolla, CA 92037. These authors contributed equally to this work (Santiago Pineda, Vesna Nikolova-Krstevski, Christiana Leimena). K. Ocorr and D. Fatkin are joint senior authors.

## Abstract

**Background:** *KCNMA1* encodes the α-subunit of the large-conductance Ca^2+^-activated K^+^ channel, K_Ca_1.1, and lies within a linkage interval for atrial fibrillation (AF). Insights into the cardiac functions of K_Ca_1.1 are limited and *KCNMA1* has not been investigated as an AF candidate gene.

**Methods and Results:** *KCNMA1* sequencing in 118 patients with familial AF identified a novel complex variant in one kindred. To evaluate potential disease mechanisms, we first evaluated the distribution of K_Ca_1.1 in normal hearts using immunostaining and immunogold electron microscopy. K_Ca_1.1 was seen throughout the atria and ventricles in humans and mice, with strong expression in the sinus node. In an *ex vivo* murine sinoatrial node preparation, addition of the K_Ca_1.1 antagonist, paxilline, blunted the increase in beating rate induced by adrenergic receptor stimulation. Knockdown of the K_Ca_1.1 ortholog, *kcnma1b*, in zebrafish embryos resulted in sinus bradycardia with dilatation and reduced contraction of the atrium and ventricle. Genetic inactivation of the *Drosophila* K_Ca_1.1 ortholog, *slo*, systemically or in adult stages, also slowed the heartbeat and produced cardiac arrhythmias.

Electrophysiological characterization of *slo-*deficient flies revealed bursts of action potentials, reflecting increased events of fibrillatory arrhythmias. Flies with cardiac-specific overexpression of the human *KCNMA1* mutant also showed increased heart period and bursts of action potentials, similar to the K_Ca_1.1 loss-of-function models.

**Conclusions:** Our data point to a highly conserved role of K_Ca_1.1 in sinus node function in humans, mice, zebrafish and fly and suggest that K_Ca_1.1 loss of function may predispose to AF.

---

Atrial fibrillation (AF) is the most common cardiac arrhythmia and a major cause of morbidity and mortality. Genetic factors were first implicated in AF pathogenesis over two decades ago when a landmark study identified a novel locus on chromosome 10q22-q24 in 3 kindreds.^1^ Two additional cardiac disorders (dilated cardiomyopathy, hypoplastic left heart syndrome) and three neurological disorders were subsequently mapped to the same chromosomal region (Figure I in the Data Supplement).^2-6^ *KCNMA1*, a gene that encodes the α-subunit of the large-conductance calcium (Ca^2+^)-activated potassium (K^+^) channel, K_Ca_1.1, lies within all 6 linkage intervals. *KCNMA1* variants have been identified in patients with neurological defects^4^ but early studies failed to detect *KCNMA1* transcripts in the heart^7,8^ and it has not been considered as a candidate gene for the mapped cardiac disorders.

K_Ca_1.1 channels are comprised of four pore-forming α-subunits that each contain a short extracellular N-terminus, 7 transmembrane domains including pore-gate and voltage-sensing domains, and a long intracellular C-terminal tail with high affinity binding sites for Ca^2+^ and multiple ligands (Figure 1). K_Ca_1.1 activation results in K^+^ efflux and membrane hyperpolarization that leads to a reduction of L-type Ca^2+^ channel activity and Ca^2+^ influx, protecting cells from Ca^2+^overload. K_Ca_1.1 channels are sensitive to Ca^2+^ and voltage and can be modulated by interactions with β and γ subunits, alternative splicing of *KCNMA1* transcripts, post-translational modification and exogenous factors.^9,10^ They contribute to Ca^2+^ homeostasis in multiple cell types and have been implicated in diverse processes, including neurotransmission, hearing, circadian rhythms, bladder control, vascular smooth muscle tone, obesity and cancer.^9,10^

**Figure 1.**
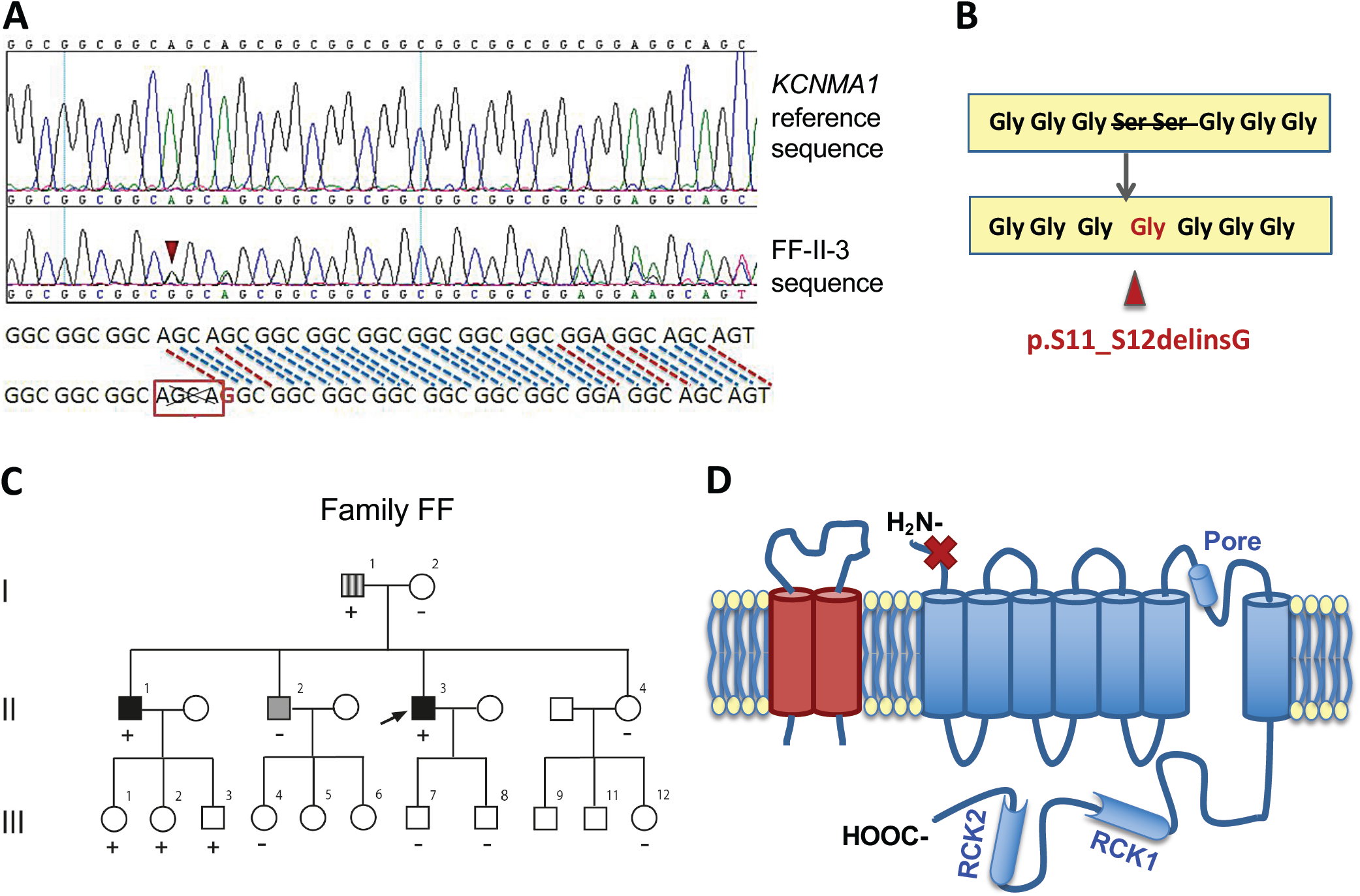
*KCNMA1* sequence variant identified in a family with tachy-brady syndrome. **(A)** *KCNMA1* sequence traces showing variation identified in the Family FF proband, FF-II-3. (**B**) This sequence change (AGCA deletion, G insertion) results in loss of two serines at amino acid positions 11 and 12, with the addition of a glycine. (**C**) Family pedigree, with phenotypes denoted as: tachy-brady syndrome (solid symbols), suspected AF (hashed symbol), no AF (open symbols); FF-II-2 had a single episode of AF associated with myopericarditis (gray symbol). The proband is indicated by an arrow. The presence (+) or absence (-) of the p.S11_S12delinsG variant are indicated. (**D**) Schematic showing K_Ca_1.1 (α-subunit, blue) with its pore and two RCK (regulator of potassium conductance) domains and a regulatory β-subunit (pink). The location of the p.S11_S12delinsG variant in the short extracellular amino terminus of the α-subunit is indicated (red cross).

The role of K_Ca_1.1 in the heart is incompletely understood. Cardiac K_Ca_1.1 channels have mainly been studied with respect to mitochondrial properties and protection against ischemia-reperfusion injury.^11-13^ K_Ca_1.1 expression has been seen in cardiac fibroblasts, telocytes, coronary artery smooth muscle and intracardiac neurons, suggesting pleiotropic functional effects.^14-17^ In recent years, there has also been accumulating evidence that K_Ca_1.1 channels are involved in heart rate regulation.^13,18-20^ Given established links between sinus node function and atrial arrhythmogenesis,^21^ these findings re-focus interest on *KCNMA1* as a potential candidate gene for AF. Here we report genetic screening of *KCNMA1* in a cohort of patients with familial AF. We evaluated K_Ca_1.1 distribution and function in the hearts of humans, mice, zebrafish, and fly, and a novel *KCNMA1* mutation was functionally characterized.

## METHODS

An expanded Methods section is provided in the Supplemental Material.

### Study Subjects

The study population was comprised of 118 individuals (79 males), aged 18 to 90 years (mean 56) with suspected familial AF, defined by AF in 2 or more first-degree relatives. Study subjects were evaluated by history and physical examination, ECG and transthoracic echocardiography. All participants were of self-reported European ancestry and provided informed written consent. Protocols were approved by St Vincent’s Hospital Human Research Ethics Committee.

### *KCNMA1* Sequencing

Protein-coding sequences and adjacent intronic sequences of the *KCNMA1* gene were PCR-amplified from genomic DNA and evaluated using Sanger sequencing. A novel *KCNMA1* variant was investigated in one family using Sanger sequencing and restriction enzyme digestion. The minor allele frequency of identified variants was evaluated in the Genome Aggregation Database (gnomAD v3; accessed June 2020).

### Human and Murine Tissue Studies

Right atrial and right ventricular tissue samples were obtained from non-failing human hearts and from adult wild-type mice. The sinoatrial node region was identified as described.^22,23^ The distribution of K_Ca_1.1 protein in tissue sections, whole mounts of the sinoatrial node complex and in isolated cardiomyocytes was evaluated by immunofluorescence microscopy using rabbit polyclonal anti-K_Ca_1.1 antibodies, and mouse monoclonal anti-ryanodine receptor-2 (RyR2), anti-caveolin-3, anti-connexin-43, and anti-Ca_v_1.3 antibodies.

For immunogold electron microscopy, fixed tissues were immunolabelled using primary rabbit polyclonal anti-K_Ca_1.1 antibodies and secondary goat anti-rabbit IgG conjugated with 15 nm gold particles. Double labelling of K_Ca_1.1 with RyR2 or Ca_v_1.2 was performed using mouse monoclonal anti-RyR or anti-Ca_v_1.2 antibodies, together with secondary goat anti-rabbit IgG conjugated with 15 nm gold particles and protein A/G conjugated with 10 nm gold particles, respectively. Sections were stained with uranyl acetate and lead citrate and examined by a JEM-1400 transmission electron microscope.

Total RNA was isolated from tissue samples and K_Ca_1.1 transcript levels were evaluated using quantitative real-time PCR (qPCR). Total protein extracts were subjected to Western blotting using rabbit polyclonal anti-K_Ca_1.1 antibodies. Blots were normalized to β-tubulin and hybridization signals were quantified with ImageJ software.

### Sinoatrial Node Ca^2+^ Imaging

Murine sinoatrial nodes were dissected, stained with a fluorogenic Ca^2+^-sensitive dye, Cal-520, bathed in oxygenated warm Tyrode’s solution, and viewed with a wide field microscope. The periodicity of Ca^2+^ transients at baseline and following addition of the K_Ca_1.1 antagonist, paxilline, ± isoproterenol, were determined.

### Zebrafish Studies

Wild-type TE zebrafish were maintained under standard aquarium conditions. Expression of zebrafish K_Ca_1.1 orthologues, *kcnma1a* and *kcnma1b*,^24^ was evaluated by reverse transcription-polymerase chain reaction using RNA isolated from whole embryo and heart extracts. Morpholino antisense oligonucleotides targeting the translation start sites of *kcnma1a* and *kcnma1b*, or a standard control morpholino, were diluted to 150μM and 2 nL then injected into 1-2 cell embryos, with the efficacy of knockdown confirmed by Western blot analysis of whole embryo lysates. Cardiac function was assessed at 3 days post fertilization (dpf). Heart rate was measured by direct microscopic observation while atrial and ventricular size and contractile function were measured by video analysis.^25^

### *Drosophila* Studies

Fly lines with systemic and cardiac-specific knockdown of the *Drosophila* K_Ca_1.1 orthologue, *slowpoke* (*slo*) were studied. To achieve *slo* knockdown in adult hearts, the Hand GeneSwitch system (HandGS-Gal4) was used.^26^ Administration of mifepristone (RU486, 100ug/mL, dissolved in ethanol) induces the HandGS-Gal4 driver, allowing temporally controlled cardiac-specific expression of *slo* RNAi. PBac *slo* genomic rescue flies were generated by injecting a *slo* genomic BAC library (gift of Hugo Bellen) into wild-type flies. Recording of semi-intact hearts was performed as described^27^ and quantified using the semi-automated optical heartbeat analysis software.^28^ For electrophysiological evaluation, semi-intact heart preparations were incubated in artificial hemolymph containing 10 µM blebbistatin (to block active crossbridge cycling). Fresh saline without blebbistatin was added and electrical potentials were recorded using sharp glass electrodes (20-50 MΩ) filled with 3M KCl using standard techniques.

### Statistical Analysis

Differences between groups were assessed using Student’s t test, Mann-Whitney test or ANOVA. Data are expressed as mean ± SEM. A p value <0.05 was considered statistically significant.

## RESULTS

### Identification of *KCNMA1* Sequence Variants

The *KCNMA1* gene was evaluated in 118 probands with familial AF. Six coding sequence variants were identified, including 4 synonymous variants and a 3-nucleotide insertion that did not alter the reading frame (Table VI in the Data Supplement). A complex variation comprised of a 4-nucleotide deletion and a 1-nucleotide insertion was identified in the proband of Family FF, II-3 (Figure 1). This sequence change, p.S11_S12delinsG, results in loss of an MspA1I enzyme restriction site, and was independently confirmed in the proband and evaluated in family members using both sequencing and restriction enzyme digestion (Figure VI in the Data Supplement). It was present in all affected family members as well as in three asymptomatic children and was absent in older unaffected family members and >70,000 unrelated subjects in the gnomAD population database. The proband had previously undergone genetic screening of 14 AF-associated K^+^ channel genes^25,29^ as well as the *SCN5A, NPPA, GJA1, GJA5, GJA7, HCN1, HCN2*, and *HCN4* genes, with no other pathogenic variants found.

FF-II-3 was diagnosed with paroxysmal lone AF at 34 years of age. He subsequently developed features of tachy-brady syndrome, with sinus bradycardia, frequent sinus pauses (up to 3 s) and permanent AF (Table VII in the Data Supplement). His affected brother, FF-II-1, had sinus node dysfunction that required pacemaker insertion and permanent AF. Their father had evidence of bradycardia on a 24-hour Holter monitor, with frequent palpitations but no documented AF. To investigate how this *KCNMA1* variant might predispose to sinus node dysfunction and AF, we next undertook a series of experiments to evaluate K_Ca_1.1 channels in normal cardiac structure and function prior to evaluating variant effects.

### Cardiac Expression of K_Ca_1.1 in Humans and Mice

The subcellular localization of K_Ca_1.1 in cardiac tissues was evaluated using immunostaining. In human right atrial tissue and isolated atrial cardiomyocytes from adult wild-type mice, K_Ca_1.1 showed a weak striation pattern with relatively stronger expression in the intercalated discs and sarcolemma (Figure 2A). K_Ca_1.1 staining overlapped with RyR2 and connexin-43 (Figure 2A). Similar patterns were observed in the human right ventricle (Figure II in the Data Supplement). In humans and in mice, there was robust expression of K_Ca_1.1 in the sinoatrial node where its expression overlapped considerably with RyR2 and the L-type calcium channel, Ca_v_1.3 (Figure 2A-C). Further evaluation of K_Ca_1.1 in human right atrial tissue was undertaken using immunogold electron microscopy (Figure 3). Gold-labelled K_Ca_1.1 epitopes were abundant in the cardiomyocyte T-tubules and sarcoplasmic reticulum, and were also present in the nucleus, sarcolemma, intercalated discs, mitochondria, coronary vascular endothelium, and fibroblasts. Negative controls for anti K_Ca_1.1 APC-021 antibody are shown in Figure IV in the Data Supplement.

**Figure 2.**
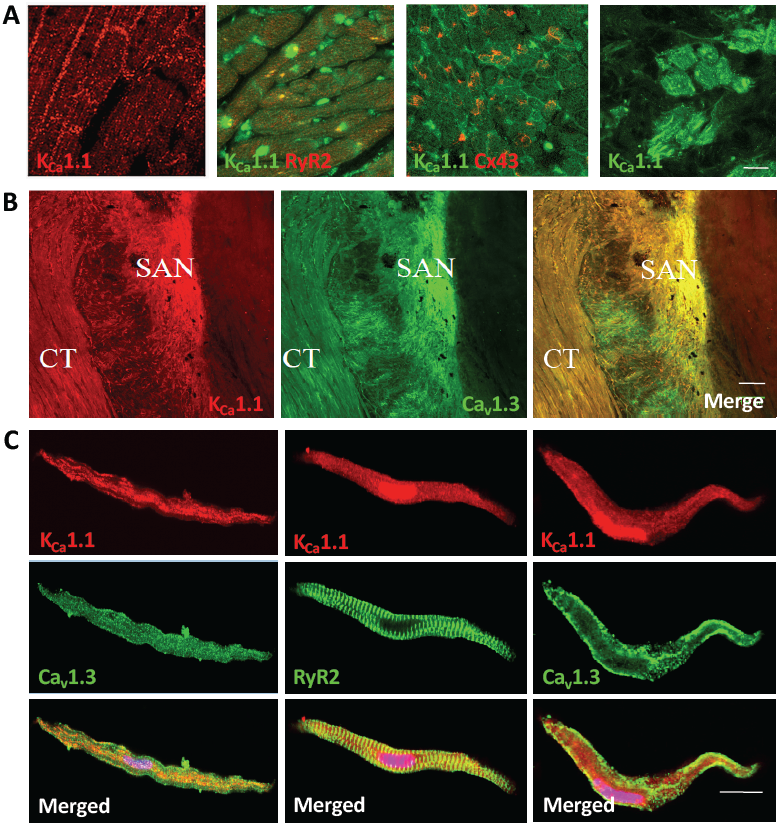
K_Ca_1.1 in the atrium. (**A**) Immunostaining of human right atrial tissue sections shows co-localization of K_Ca_1.1 with the ryanodine receptor (RyR2) and connexin-43 (Cx43), with strong K_Ca_1.1 expression in the sinus node (right panel); scale bar = 5 μm. (**B**) Whole mount immunostaining of the murine sinoatrial node (SAN) complex. K_Ca_1.1 is expressed throughout the SAN and paranodal connective tissue (CT) and co-localizes with the L-type calcium channel, Ca_v_1.3; scale bar = 100 μm. (**C**) Isolated murine atrial cardiomyocytes (left column) and pacemaker cells (center and right columns) show K_Ca_1.1 co-localization with RyR2 and Ca_v_1.3; scale bar = 10 μm. Antibody optimization and a negative control for immunostaining are shown in Figures II and III in the Data Supplement.

**Figure 3.**
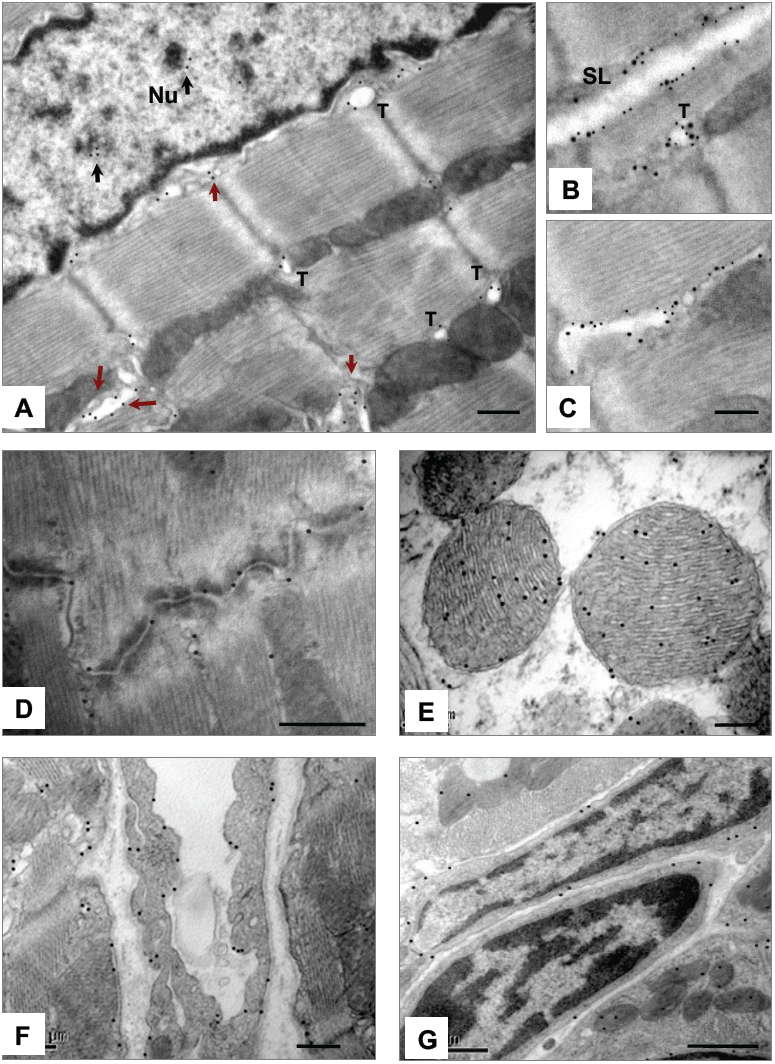
Immunogold electron microscopy of K_Ca_1.1 in human atrial tissue. (**A**) In atrial cardiomyocytes, colloidal gold labelled K_Ca_1.1 (black dots) was present in the nucleus (nu), T-tubules (T), and sarcoplasmic reticulum (red arrows). Double-labelling of K_Ca_1.1 (15 nm particles, large black dots) and L-type calcium channel, Ca_v_1.2 or ryanodine receptor-2 (10 nm particles, small block dots), shows K_Ca_1.1 in (**B**) the sarcolemma (SL) and T-tubules (T), and (**C)** sarcoplasmic reticulum. K_Ca_1.1 was also present in (**D**) intercalated discs, (**E**), mitochondria, (**F**) coronary vascular endothelium and (**G**) fibroblasts. Negative controls for immunolabelling are shown in Figure IV in the Data Supplement. Scale bars = (**A**,**D**,**E**) 0.2 μm; (**B, C**) 0.1 μm; (**F**) 0.5 μm; (**G**) 1 μm.

The presence of *KCNMA1* transcript in human right atrial tissue was confirmed by qPCR with higher levels of expression in the sinus node than in paranodal regions and in atrial myocardium (Figure VIII in the Data Supplement). Total *KCNMA1* transcript levels were higher in atrial tissue from patients with a history with AF when compared to those without AF (Figure 4). These differences appeared to be driven by the subset of older patients (≥70 years) who had atrial dilatation and chronic AF (Figure 4B-C; Figure IX in the Data Supplement). We then evaluated expression of a number of *KCNMA1* isoforms (Figure 4C). Levels of the mitochondrial-specific isoform, DEC, were also significantly higher in older patients with AF when compared to age-matched patients without AF and younger patients with AF. Levels of the stretch-activated isoform, STREX, increased with age in patients with and without AF (Figure 4C). On Western blotting, two K_Ca_1.1 bands were detected that were approximately 100 kD and 50 kD, and K_Ca_1.1 protein levels were increased in patients with AF (Figure 4D-E).

**Figure 4.**
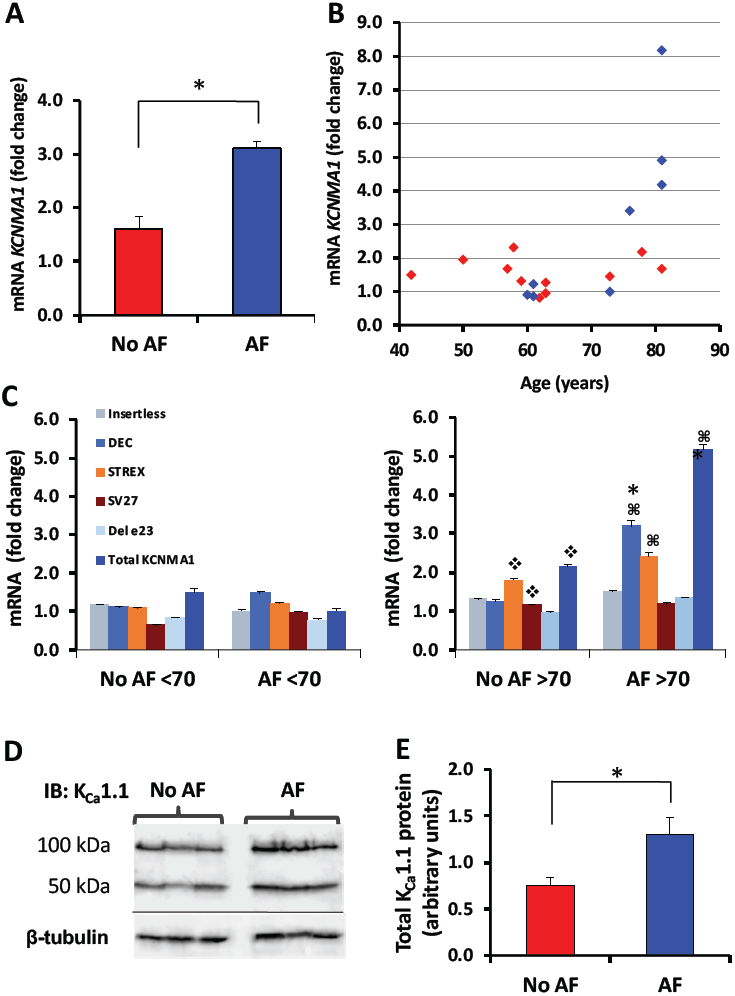
Quantititive assessment of atrial K_Ca_1.1 expression. (**A**) qPCR evaluation of *KCNMA1* mRNA in right atrial appendage tissue from patients in sinus rhythm with normal atrial size (red, n=11) and patients with atrial dilatation and documented AF (blue, n=8). (**B**) Dot blot representation of total *KCNMA1* mRNA levels in individual patients (from **A**) showing effects of AF status and age. (**C**) Mean data for mRNA levels of total *KCNMA1* and five isoforms in patients with and without a history of AF, aged <70 years or ≥70 years at the time of study; ❖*P*<0.05 vs no AF <70; □ *P*<0.05 vs AF <70; * *P*<0.05 vs no AF ≥70. (**D**) Western blot showing total K_Ca_1.1 protein in patients with and without AF (n=3 in each group); β-tubulin was used as a loading control. Negative controls for Western blotting are shown in Figure V in the Data Supplement. (**E**) Mean data for levels of total K_Ca_1.1 protein in patients with (n=8) and without AF (n=11); * *P*<0.05.

### Effects of Adrenergic Receptor Stimulation and Paxilline on Sinoatrial Node Function

To investigate the role of K_Ca_1.1 in sinoatrial node function, we made use of an *ex vivo* preparation of the murine sinoatrial node (Figure 5A). Addition of the K_Ca_1.1 antagonist, paxilline 10 µM, alone did not significantly reduce the beating rate (control: 359 ± 33; paxilline: 321 ± 18, *P*=0.36; n=8). When the sinoatrial node was subjected to an adrenergic stimulus, isoproterenol 500 nM, the Ca^2+^ transient frequency increased from 340 ± 23 beats per minute (bpm) to 520 ± 19 bpm (*P*=0.0003; Figure 5B). This adrenergic-induced rate increase was blunted in the presence of paxilline 10 µM, with an average of 440 ± 18 bpm (*P*=0.0.03; n=6; Figure 5B-F), suggesting K_Ca_1.1-dependent effects.

**Figure 5.**
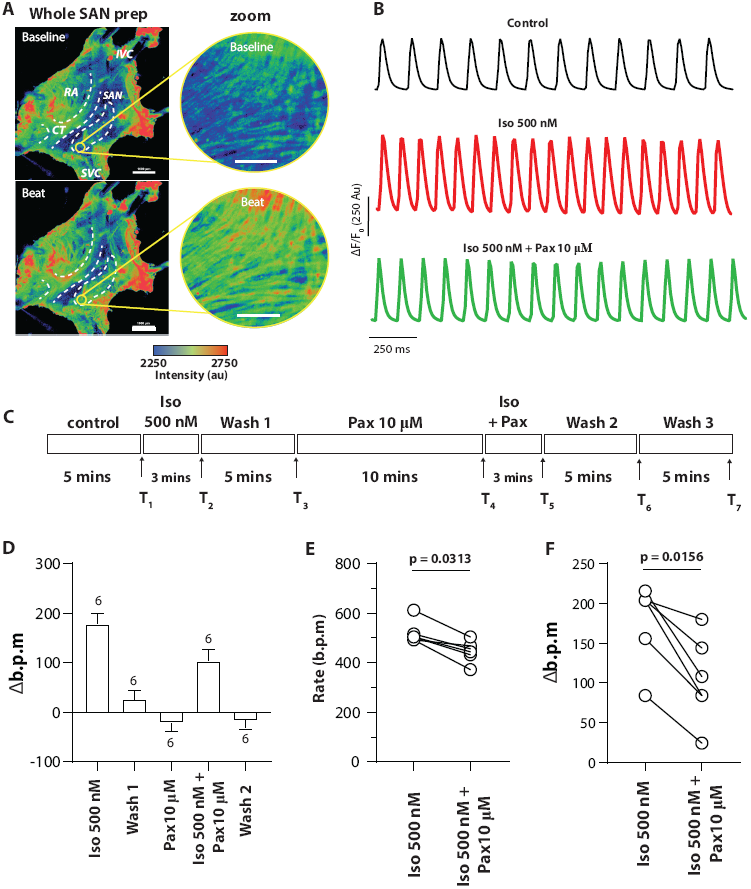
Effect of K_Ca1.1_ blockade on *ex vivo* murine sinoatrial node (SAN) function. (**A**) Images of Cal520-loaded whole SAN preparation showing the right atrium (RA), crista terminalis (CT), inferior vena cava (IVC) and superior vena cava (SVC). Intensity of the intracellular Ca^2+^ signal is shown at baseline (between beats [top panel], zoom [right]) and at the peak of a Ca^2+^ transient (during beat [lower panel], zoom [right]). (**B**) Raw traces of Ca^2+^ transients from the Ca^2+^ signal in arbitrary fluorescent units (au) for control, isoproterenol 500nM, and isoproterenol 500nM plus paxilline 10 µM. (**C**) Timeline showing the experimental protocol with incubation times. A 5 s imaging measurement was taken at each of 7 time points (T1-T7). (**D**) Effects of isoproterenol 500 nM (Iso) and paxilline 10 µM (Pax) shown as change in beating rate (Δ b.p.m) compared to control recordings (n=6; data shown as mean ± SEM). (**E)** Difference in absolute beating rate for SAN treated with isoproterenol 500 nM alone or isoproterenol 500 nM plus paxilline 10 µM (n=6). (**F**) Difference in Δ b.p.m between SAN treated with isoproterenol 500 nM alone and isoproterenol 500 nM plus paxilline 10 µM (n=6).

### K_Ca_1.1 Knockdown Alters Cardiac Function in Zebrafish

The zebrafish orthologous genes, *kcnma1a* and *kcnma1b*, share ∼90% sequence identity and have ∼85% homology to human *KCNMA1*.^24^ Both *kcnma1* genes were expressed in zebrafish whole embryos and embryonic heart at 3 dpf (Figure 6A). Morpholinos were designed to target the translation start sites of the *kcnma1a* and *kcnma1b* genes with effective protein depletion achieved for *kcnma1b* (Figure 6B; Figure X in the Data Supplement). Overall size and morphology of the *kcnma1b* morphant was grossly normal. *Kcnma1b* morpholino-injected embryos had lower mean heart rate when compared to control-injected embryos (123 ± 4 bpm vs 140 ± 2 bpm, *P*<0.001), with larger atrial diameter (108 ± 5 μm vs 79 ± 3 μm, *P*<0.0001) and reduced atrial fractional area change (27 ± 2% vs. 47 ± 2%, *P*<0.0001) (Figure 6C-D). Ventricular end-diastolic diameter (*P*<0.0001) and fractional shortening (*P*<0.0001) were lower in *kcnma1b* morpholino-injected embryos than in controls (Figure XI in the Data Supplement). These observations indicate that *kcnma1b* depletion affects heart rate, as well as atrial and ventricular chamber formation and function in zebrafish.

**Figure 6.**
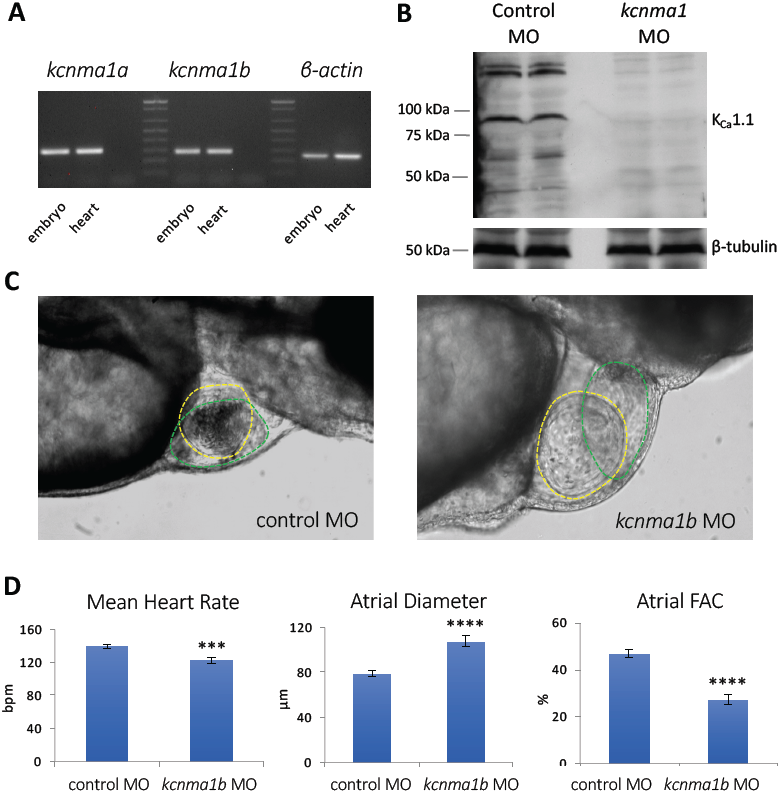
Characterization of zebrafish *kcnma1*. (**A**) Expression of *kcnma1* genes in whole embryo and isolated heart of 3 dpf wild-type zebrafish, detected by RT-PCR; β-actin was used as a control for cDNA quality. (**B**) Western blot showing *kcnma1* protein levels in control morpholino (MO)-injected and *kcnma1b* morpholino-injected embryos; samples were run in duplicate, β-tubulin was used as a loading control. Western blot was performed with anti-K_Ca_1.1 antibody specific for the *kcnma1b* isoform. (**C**) Brightfield images of hearts of 3 dpf zebrafish embryos injected with either control or *kcnma1b* morpholino showing relative sizes of the atria (yellow dashed line) and ventricle (green dashed line). (**D**) Mean data for heart rate, maximal atrial diameter and atrial fractional area change (FAC) in control morpholino-injected and *kcnma1b* morpholino-injected embryos at 3 dpf (n=17-20 each group); **** *P*<0.0001, *** *P*<0.001.

### K_Ca_1.1 is a Determinant of Cardiac Function in *Drosophila*

The *Drosophila slowpoke* (*slo*) gene has ∼60% sequence identity to the human *KCNMA1* gene and is reportedly present in the fly heart.^30^ We confirmed the cardiac expression of *slo* using nanofluidic qPCR (Figure XII in the Data Supplement). To determine the effects of K_Ca_1.1 deficiency in the *Drosophila* model, we reduced systemic or cardiac *slo* function and evaluated cardiac parameters using semi-automated optical heartbeat analysis^27,28^ in young (1 week), middle-aged (3 week) and old (5 week) mutants and age-matched controls.

We first generated flies with two loss-of-function *slo* alleles: one of these, *slo*^*4*^, is a null allele produced by a gamma ray-induced chromosomal inversion,^31^ and the other, Df(3R) BSC 397, contains a transposon-mediated deletion. These trans-heterozygote flies (slo^4^/Df(3R) BSC 397) showed increased heart period and a higher arrhythmia index (defined as irregularly irregular heart period)^27^ at 3 weeks of age when compared to control flies (Figure 7A-B; Figure XII in the Data Supplement). To show that *slo* deficiency is the cause of these heart phenotypes, we crossed a wild-type genomic copy of *slo* into the slo^4^/Df(3R) BSC 397 combination. In these flies, the slow heartbeat and arrhythmic phenotype were rescued (Figure 7A-B). Interestingly, the presence of one or two extra copies of genomic *slo* in the wild-type fly background also markedly increased the heart period (Figure XIII A-D in the Data Supplement). Taken together, these results show that heart rate regulation in flies is sensitive to K_Ca_1.1 dose, with both deficiency and excess resulting in bradycardia.

**Figure 7.**
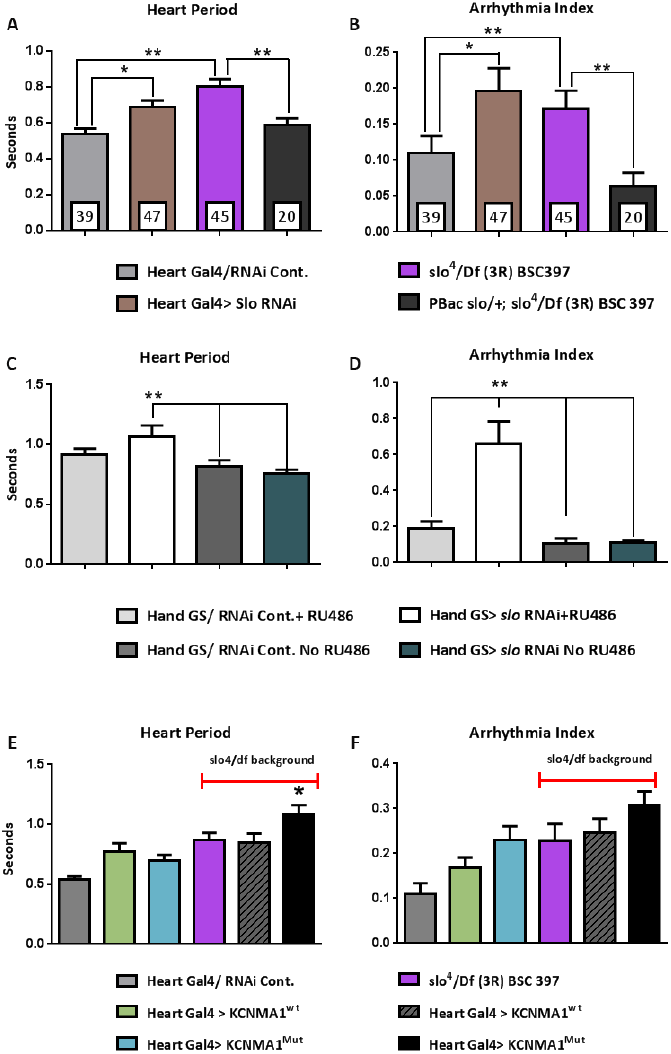
Cardiac function in *Drosophila slo* mutants at 3 weeks of age. (**A**) Heart period and (**B**) arrhythmia index are increased in flies with cardiac-specific RNAi knockdown of *slo* (Heart Gal4>Slo RNAi) and a trans-heterozygous configuration of two mutant *slo* alleles (slo^4^/Df(3R) BSC 397) compared to controls. An extra copy of the wild-type *slo* genomic locus (PBac slo/+; slo^4^/Df(3R) BSC 397) was able to rescue the cardiac phenotype of the transheterozygous mutant. (**C**,**D**) Cardiac-specific *slo* knockdown in adult flies resulted in a dramatic increase in arrhythmia index compared to controls. (**E**,**F**) Cardiac expression of the human p.S11_S12delinsG *KCNMA1*, but not the wild-type form, resulted in a significantly increased heart period (**E**) with a trend towards a higher arrhythmia index (**F**) in a sensitized background. For all genotypes, n=20-50 each group. \**P*<0.05; \*\**P*<0.01.

To determine whether *slo* acted directly in the heart, we crossed the cardiac Gal4 driver, Hand4.2-Gal4, and UAS-*slo*-RNAi lines to generate flies with heart-specific knockdown of *slo* (“Heart Gal4>slo RNAi”). When compared to controls, the Heart Gal4>slo RNAi flies, like systemic *slo*-deficient flies, had significant prolongation of diastolic and systolic intervals, resulting in an overall increase in the heart period (slower heart rate) and increased arrhythmias at 3 weeks of age (Figure 7A-B; Figure XII A-D in the Data Supplement). Similar changes were seen at 5 weeks, although for arrhythmia index, the control values increased with age as expected^27^ and the differences between groups were not statistically significant (Figure XII A-D in the Data Supplement).

The Heart Gal4>slo RNAi flies have *slo* knockdown in the heart throughout development and a significant proportion died during aging (Figure XII E in the Data Supplement). To determine if *slo* knockdown during development or at adult stages was important, we used a conditional heart driver, HandGS-Gal4 to restrict knockdown to adult stages.^26^ Adult-only *slo* knockdown in the heart also caused a slower heartbeat and substantial arrhythmias (Figure 7C-D; Figure XIV in the Data Supplement). These data suggest that adult-only *slo* knockdown is sufficient to cause cardiac abnormalities.

To exclude the possibility that neural *slo* function might be responsible for this cardiac phenotype, we crossed a pan-neural Gal4 driver line (elav-Gal4) with the UAS-slo-RNAi line. We found no significant differences in either heart period or arrhythmia index points between mutant (“Neural Gal4>slo RNAi”) and control flies (Figure XIII E-F in the Data Supplement), suggesting a cardiac autonomous effect of *slo* deficiency.

Electrophysiological evaluation of fly hearts was performed at 3 weeks of age. Intracellular tracings in wild-type (w^1118^) and in Heart Gal4/RNAi control hearts typically showed a resting membrane potential of -40 to -60mV, with single action potential peaks of 60 mV amplitude (Figure 8; Table VIII in the Data Supplement). There were no changes in resting membrane potential, maximum action potential amplitude, or time to maximum action potential amplitude in the *slo* mutants when compared to wild-type flies. However, both Heart Gal4>slo RNAi and slo^4^/Df(3R) BSC397 flies showed multiple peaks and a longer duration of depolarization in each action potential suggestive of diminished repolarization reserve and increased propensity to early after-depolarizations. In particular, flies with conditional adult-only *slo* knockdown showed frequent busts of action potentials (Figure 8E), consistent with the high incidence of arrhythmias evident in M-mode tracings (Figure XIV in the Data Supplement).

**Figure 8.**
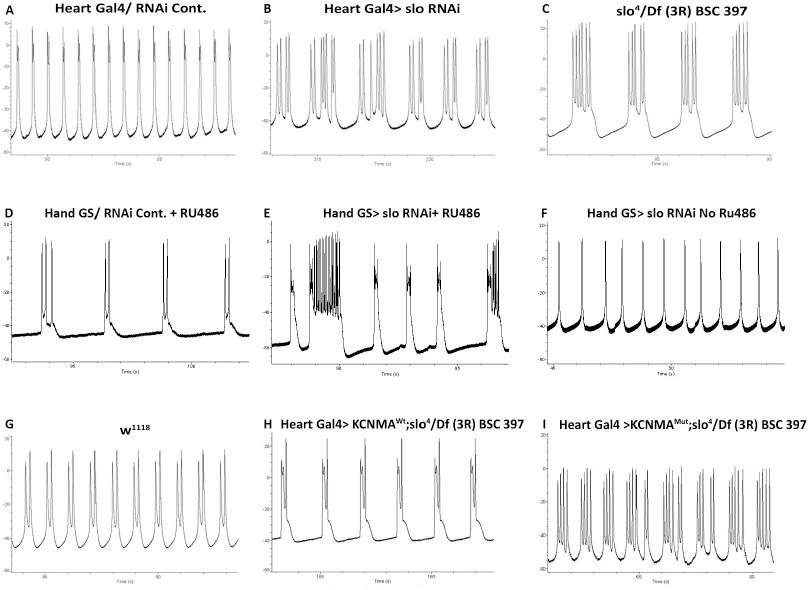
Electrophysiological evaluation of *Drosophila slo* mutants. Ten second representative traces of the electrical activity within control lines: (**A, D, F, G)**, *slo-*deficient lines: (**B, C, E)** and human transgene expression in *slo* background (**H**,**I**); *slo* knockdown lines showed increased peaks per event and higher event duration compared to controls (see Table VIII in the Data Supplement); Y axis: voltage (mV), X axis: time (ms). Cardiac-specific human wild-type *KCNMA1* expression showed a prolonged contraction phenotype but this was less pronounced than seen with the human *KCNMA1* mutant in a null background.

### Functional Characterization of Human *KCNMA1* Mutation

To study the p.S11_S12delinsG mutation identified in Family FF, wild-type and mutant human *KCNMA1* constructs were generated and injected into w^1118^ flies as UAS transgenes, then crossed into the *slo*^*4*^ mutant background. This strategy created a sensitized background to evaluate the human *KCNMA1* constructs in the absence of the endogenous *slo* gene. At 3 weeks of age, there were no differences in heart period, arrhythmia index or action potential characteristics between flies with cardiac-specific overexpression of wild-type human *KCNMA1* (“Heart Gal4>KCNMA1^wt^”) compared to *slo* mutants alone (Figure 7E-F and Figure 8H). In contrast, flies overexpressing the human *KCNMA1* mutant (“Heart Gal4>KCNMA1^mut^”) showed a significantly longer heart period (*P*<0.05), a trend towards increased arrhythmia index (*P*=0.05), and had relatively longer action potential burst events and a greater number of peaks per burst when compared to *slo* control groups (Figure 7E-F and Figure 8I). Collectively, these results indicate that wild-type human *KCNMA1* is unable to rescue flies with *slo* deficiency, but mutant human *KCNMA1* is functionally deleterious and exacerbates the effects of *slo* deficiency.

## DISCUSSION

Our data provide new perspectives on the role of K_Ca_1.1 in the normal and diseased heart and highlight its role in sinus node function. Although early work in *Drosophila* larvae suggested that K_Ca_1.1 was involved in heart rate regulation,^32^ more recent studies in mice have rekindled interest in these channels as potential determinants of cardiac pacemaker function. Imlach et al.^18^ noted that resting heart rates in *Kcnma*-/- and wild-type mice were similar.

Pharmacological inhibition of K_Ca_1.1 using paxilline and iberiotoxin reduced heart rates in wild-type mice and in isolated rat hearts but had no effects in homozygous *Kcnma1* knockout (*Kcnma*-/-) mice. Subsequent analysis by Lai et al.^19^ showed that baseline firing rates of isolated sinus node cells from *Kcnma*-/- mice were reduced when compared to wild-type cells, and it was proposed that the absence of bradycardia in intact animals was due to compensatory activation of the sympathetic nervous system. Modest reductions in heart rate, ventricular ejection fraction and mean arterial pressure were seen by Frankenreiter et al.^13^ in mice with cardiomyocyte-specific K_Ca_1.1 deletion. In another *in vivo* model, Patel et al.^20^ reported that injection of paxilline in wild-type rats elicited dose-dependent reductions of heart rate and coronary flow. Here we show that K_Ca_1.1 is highly expressed in the sinus node in humans as well as in mice, and that K_Ca_1.1 inactivation affects heart rate in mice, embryonic zebrafish and adult flies. Extra copies of K_Ca_1.1 in the fly had similar effects, indicating that a narrow range of K_Ca_1.1 levels is required to maintain normal sinus node function.

Collectively these data raise intriguing questions about how K_Ca_1.1 contributes to heart rate regulation and arrhythmogenesis. In the “Ca^2+^ clock” model of pacemaker automaticity, impulse generation is dependent on cyclical RyR2-mediated Ca^2+^ release from sarcoplasmic reticulum stores coupled to Ca^2+^ inflow though voltage-gated Ca^2+^channels.^33^ The latter include L-type Ca_v_1.3 channels that are activated at negative voltages at the beginning of diastolic depolarization. Our finding of K_Ca_1.1 in the vicinity of RyR2 and Ca_v_1.3 in pacemaker cells suggests that activation of K_Ca_1.1 might be a key component of this cycle, linking Ca^2+^ release, membrane hyperpolarization and Ca_v_1.3 activation. In this model, K_Ca_1.1 deficiency would be anticipated to prolong diastolic depolarization by limiting intracellular Ca^2+^ repletion. It is notable that bradycardia is a phenotypic feature of human mutations and animal models that have excessive RyR2 activity or reduced Ca_v_1.3 activity, respectively.^34-37^ Previous studies have shown that K_Ca_1.1 is highly expressed in fibroblasts, which are particularly abundant in the sinus node.^38^ Interactions between fibroblasts and myocytes are an important component of mechano-electric feedback in the atrium and can influence the spontaneous depolarization rates of pacemaker cells. Hence, K_Ca_1.1 deficiency could also impact on fibroblast-myocyte coupling with effects on basal heart rate and on chronotropic responses to atrial stretch.

There is increasing recognition that the combination of bradycardia and AF (“tachy-brady” syndrome) may be indicative of a diffuse process that extends beyond the sinus node to involve atrial myocardium.^33,39^ Here we find that K_Ca_1.1 has an extensive subcellular distribution in human atrial cardiomyocytes, as well as in fibroblasts and coronary vessels. K_Ca_1.1 knockdown resulted in cardiac chamber dilatation and impaired contractile function in zebrafish embryos, with prolonged action potential duration and increased action potential spikes in flies. We previously reported that K_Ca_1.1 is a downstream effector of interactions between the Rho-GTPase *Cdc42* and the cardiac transcription factor *Nkx2*.*5*. Compound heterozygous *Cdc42*-*Nkx2*.*5* mutant flies and mice showed reduced K_Ca_1.1 expression and changes in myofibrillar architecture, cardiac output, and conduction.^30^ Taken together with work by others, these findings point to numerous ways in which K_Ca_1.1 deficiency could contribute to an atrial substrate for arrhythmogenesis, including atrial electrical and structural remodelling, myocardial ischemia, oxidative stress, and altered autonomic tone.

For the human *KCNMA1* mutation identified in Family FF, the segregation analysis was consistent with disease association, and evaluation in flies suggested a K_Ca_1.1 loss of function effect. The p.S11_S12delinsG variation is located in the extracellular N-terminus of K_Ca_1.1 that is required for interactions with the channel’s β-subunits^40^ and the S12 residue has been demonstrated to be an *in vivo* K_Ca_1.1 phosphorylation site.^41^ Hence, mutation in this region could impair K_Ca_1.1 channel activation due to changes in β-subunit modulation or post-translational modification. We posit that K_Ca_1.1 loss of function is an important risk factor for AF in Family FF. Although no variants in known AF disease genes were identified, it remains possible that other deleterious genetic variants may be contributing to disease in this family. No spontaneous episodes of AF were observed in our zebrafish model but this may be due to the young age at which the fish were studied, small heart size, baseline fast heart rate, or lack of an appropriate trigger. K_Ca_1.1-deficient flies did show an increased incidence of arrhythmia, which is similar to findings in long QT models in *Drosophila* and suggestive of a deficit in repolarization capacity.

Increased levels of K_Ca_1.1 transcript in older subjects with chronic AF, when compared to young subjects with AF and age-matched individuals without AF, was an unexpected finding and most likely represents a compensatory response to mitigate against intracellular Ca^2+^ overload. This may be a contributing factor to the substantial reductions (∼70%) in atrial *I*_CaL_ that are typically seen in patients with persistent AF.^42^ While initially adaptive, increased expression of human K_Ca_1.1 has been shown to shorten the action potential duration in HL-1 atrial cardiomyocytes,^43^ and this could increase the propensity for initiation and maintenance of AF via re-entry mechanisms. The phosphodiesterase-III inhibitor, milrinone, is a heart failure therapy that has recently been shown to cause K_Ca_1.1-dependent dilatation of the pulmonary veins.^44^ AF is a recognized complication of milrinone therapy,^45^ and may be attributable, at least in part, to K_Ca_1.1 activation. In flies, we found that extra copies of the *slo* locus increased the heart period and arrhythmia index. Taken together, these findings indicate that K_Ca_1.1 excess, as well as K_Ca_1.1 deficiency can have deleterious effects on cardiac function and have implications for administration of K_Ca_1.1 activating drugs, which have been proposed as potential therapies for long QT syndrome and ischemia-reperfusion injury.^43,46^ Pharmacological manipulation of K_Ca_1.1 activity warrants investigation as a new treatment option for cardiac arrhythmias and myopathies, but as for many other cardiac ion channels, there may be a narrow beneficial therapeutic range.

Taken together, our data show that K_Ca_1.1 channels have a highly conserved role in sinus node function and arrhythmia risk. Our findings provide support for *KCNMA1* as a candidate disease gene for AF and other human cardiac disorders that map to chromosome 10q22-q24. Further studies to explore the potentially diverse roles of K_Ca_1.1 in cardiac function are likely to be fruitful.

## Acknowledgements

We thank Jamie Vandenberg for helpful discussions, and the Victor Chang Cardiac Research Institute Innovation Centre, funded by the NSW Government.

## Sources of Funding

The authors were supported by the National Health and Medical Research Council of Australia (1019693,1025008,1074386,459419,573732), National Heart Foundation of Australia (PB06S2916), Estate of the Late RT Hall, St Vincent’s Clinic Foundation, NSW Health Early Career Fellowship, Prince Charles Hospital Foundation (EN2018-01), British Heart Foundation, Fondation Leducq (TNE FANTASY 19CV03), Hunter Medical Research Institute, American Heart Association (14GRNT20490239), National Institutes of Health (HL054732, HL098053).

## Disclosures

None.

## Notes

### Competing Interest Statement

The authors have declared no competing interest.

### Summary of Updates

This manuscript contains minor edits - Figure 6 now includes a full image of the western blots and authors were added.

## REFERENCES

1. Brugada R, Tapscott T, Czernuszewicz GZ, Marian AJ, Iglesias A, Mont L, et al. Identification of a genetic locus for familial atrial fibrillation. N Engl J Med. 1997;336:905–911.

2. Bowles KR, Abraham SE, Brugada R, Zintz C, Comeaux J, Sorajja D, et al. Construction of a high-resolution physical map of the chromosome 10q22-q23 dilated cardiomyopathy locus and analysis of candidate genes. Genomics. 2000;67:109–127.

3. Hinton RB, Martin LJ, Rame-Gowda S, Tabangin ME, Cripe LH, Benson DW. Hypoplastic left heart syndrome links to chromosomes 10q and 6q and is genetically related to bicuspid aortic valve. J Am Coll Cardiol. 2009;53:1065–1071.

4. Du W, Bautista JF, Yang H, Diez-Sampedro A, You SA, Wang L, et al. Calcium-sensitive potassium channelopathy in human epilepsy and paroxysmal movement disorder. Nat Genet. 2005;37:733–738.

5. Anttila V, Nyholt DR, Kallela M, Artto V, Vepsalainen S, Jakkula E, et al. Consistently replicating locus linked to migraine on 10q22-q23. Am J Hum Genet. 2008;82:1051–1063.

6. Ratnapriya R, Satishchandra P, Kumar SD, Gadre G, Reddy R, Anand A. A locus for autosomal dominant reflex epilepsy precipitated by hot water maps at chromosome 10q21.3-q22.3. Hum Genet. 2009;125:541–549.

7. Tseng-Crank J, Foster CD, Krause JD, Mertz R, Godinot N, DiChiara TJ, et al. Cloning, expression, and distribution of functionally distinct Ca^2+^-activated K^+^ channel isoforms from human brain. Neuron. 1994;13:1315–1330.

8. Knaus HG, Eberhart A, Koch RO, Munujos P, Schmalhofer WA, Warmke JW, et al. Characterization of tissue-expressed alpha subunits of the high conductance Ca^2+^-activated K^+^ channel. J Biol Chem. 1995;270:22434–22439.

9. Contreras GF, Castillo K, Enrique N, Carrasquel-Ursulaez W, Castillo JP, Milesi V, et al. A BK (Slo1) channel journey from molecule to physiology. Channels. 2013;7:442–458.

10. Singh H, Stefani E, Toro L. Intracellular BK_Ca_ (iBK_Ca_) channels. J Physiol. 2012;590:5937–5947.

11. Xu W, Liu Y, Wang S, McDonald T, Van Eyk JE, Sidor A, et al. Cytoprotective role of Ca^2+^-activated K^+^ channels in the cardiac inner mitochondrial membrane. Science. 2002;298:1029–1033.

12. Balderas E, Zhang J, Stefani E, Toro L. Mitochondrial BK_Ca_ channel. Front Physiol. 2015;6:104.

13. Frankenreiter S, Bednarczyk P, Kneiss A, Bork NI, Straubinger J, Koprowski P, et al. cGMP-elevating compounds and ischemic conditioning provide cardioprotection against ischemic and reperfusion injury via cardiomyocyte-specific BK channels. Circulation. 2017;136:2337–2355.

14. Wang YJ, Sung RJ, Lin MW, Wu SN. Contribution of BK_Ca_-channel activity in human cardiac fibroblasts to electrical coupling of cardiomyocytes-fibroblasts. J Membrane Biol. 2006;213:175–185.

15. Sheng J, Shim W, Lu J, Lim SY, Ong BH, Lim TS, et al. Electrophysiology of human cardiac atrial and ventricular telocytes. J Cell Mol Med 2014;18:355–362.

16. Tanaka Y, Meera P, Song M, Knaus HG, Toro L. Molecular constituents of maxi K_Ca_ channels in human coronary smooth muscle: predominant α + β subunit complexes. J Physiol. 1997;502:545–557.

17. Selga E, Perez-Serra A, Moreno-Asso A, Anderson S, Thomas K, Desai M, et al. Molecular heterogeneity of large-conductance calcium-activated potassium channels in canine intracardiac ganglia. Channels (Austin). 2013;7:322–328.

18. Imlach WL, Finch SC, Miller JH, Meredith AL, Dalziel JE. A role for BK channels in heart rate regulation in rodents. PLoS One. 2010;5:e8698.

19. Lai MH, Wu Y, Gao Z, Anderson ME, Dalziel JE, Meredith AL. BK channels regulate sinoatrial node firing rate and cardiac pacing in vivo. Am J Physiol Heart Circ Physiol. 2014;307:H1327–H1338.

20. Patel NH, Johannesen J, Shah K, Goswani SK, Patel NJ, Ponnalagu D, et al. Inhibition of BK_Ca_ negatively alters cardiovascular function. Physiol Rep. 2018;6:e13748.

21. Monfredi O, Boyett MR. Sick sinus syndrome and atrial fibrillation in older persons – a view from the sinoatrial nodal myocyte. J Mol Cell Cardiol. 2015; 83:88–100.

22. Chandler NJ, Greener ID, Tellez JO, Inada S, Musa H, Molenaar P, et al. Molecular architecture of the human sinus node: insights into the function of the cardiac pacemaker. Circulation. 2009;119:1562–1575.

23. Ju YK, Chu Y, Chaulet H, Lai D, Gervasio OL, Graham RM, et al. Store-operated Ca^2+^ influx and expression of TRPC genes in mouse sinoatrial node. Circ Res. 2007;100:1605–1614.

24. Rohmann KN, Deitcher DL, Bass AH. Calcium-activated potassium (BK) channels are encoded by duplicate *slo1* genes in teleost fishes. Mol Biol Evol. 2009;26:1509–1521.

25. Liang B, Soka M, Christensen AH, Olesen MS, Larsen AP, Knop FK, et al. Genetic variation in the two-pore domain potassium channel, TASK-1, may contribute to an atrial substrate for arrhythmogenesis. J Mol Cell Cardiol. 2014;67:69–76.

26. Osterwalder T, Yoon KS, White BH, Keshishian H. A conditional tissue-specific transgene expression system using inducible GAL4. Proc Natl Acad Sci USA. 2001;98:12596–12601.

27. Ocorr K, Reeves NL, Wessells RJ, Fink M, Chen HS, Akasaka T, et al. KCNQ potassium channel mutations cause cardiac arrhythmias in Drosophila that mimic the effects of aging. Proc Natl Acad Sci USA. 2007;104:3943–3948.

28. Fink M, Callol-Massot C, Chu A, Ruiz-Lozano P, Izpisua Belmonte JC, Giles W, et al. A new method for detection and quantification of heartbeat parameters in Drosophila, zebrafish and embryonic mouse hearts. Biotechniques. 2009;46:101–113.

29. Mann SA, Otway R, Guo G, Soka M, Karlsdotter L, Trivedi G, et al. Epistatic effects of potassium channel variation on cardiac repolarization and atrial fibrillation risk. J Am Coll Cardiol. 2012;59:1017–1025.

30. Qian L, Wythe JD, Liu J, Cartry J, Vogler G, Mohapatra B, et al. R. Tinman/Nkx2-5 acts via miR-1 and upstream of Cdc42 to regulate heart function across species. J Cell Biol. 2011;193:1181–1196.

31. Atkinson NS, Robertson GA, Ganetzky B. A component of calcium-activated potassium channels encoded by the *Drosophila slo* locus. Science. 1991;253:551–555.

32. Johnson E, Ringo J, Bray N, Dowse H. Genetic and pharmacological identification of ion channels central to the *Drosophila* cardiac pacemaker. J Neurogenet. 1998;12:1–24.

33. Monfredi O, Maltsev VA, Lakatta EG. Modern concepts concerning the origin of the heartbeat. Physiology (Bethesda). 2013;28:74–92.

34. Bhuiyan ZA, van den Berg MP, van Tintelen JP, Bink-Boelkens MT, Wiesfeld AC, Alders M, et al. Expanding spectrum of human RYR2-related disease: new electrocardiographic, structural, and genetic features. Circulation. 2007;116:1569–1576.

35. Neco P, Torrente AG, Mesirca Neco P, Torrente AG, Mesirca P, Zorio E, Liu N, Priori SG, et al. Paradoxical effect of increased diastolic Ca^2+^ release and decreased sinoatrial node activity in a mouse model of catecholaminergic polymorphic ventricular tachycardia. Circulation. 2012;126:392–401.

36. Baig SM, Koschak A, Lieb A, Gebhart M, Dafinger C, Numberg G, et al. Loss of Ca_v_1.3 (*CACNA1D*) function in a human channelopathy with bradycardia and congenital deafness. Nat Neurosci. 2011;14:77–84.

37. Mangoni ME, Couette B, Bourinet E, Platzer J, Reimer D, Striessnig J, et al. Functional role of L-type Ca_v_1.3 Ca^2+^ channels in cardiac pacemaker activity. Proc Natl Acad Sci USA. 2003;100:5543–5548.

38. Yue L, Xie J, Nattel S. Molecular determinants of cardiac fibroblast electrical function and therapeutic implications for atrial fibrillation. Cardiovasc Res. 2011;89:744–753.

39. Sanders P, Morton JB, Kistler PM, Spence SJ, Davidson NC, Hussin A, et al. Electrophysiological and electroanatomic characterization of the atria in sinus node disease: evidence of diffuse atrial remodelling. Circulation. 2004;109:1514–1522.

40. Morrow JP, Zakharov SI, Liu G, Yang L, Sok AJ, Marx SO. Defining the BK channel domains required for β1-subunit modulation. Proc Natl Acad Sci USA. 2006;103:5096–5101.

41. Yan J, Olsen JV, Park KS, Li W, Bildl W, Schulte U, et al. Profiling the phospho-status of the BK_Ca_ channel α subunit in rat brain reveals unexpected patterns and complexity. Mol Cell Proteomics. 2008;7:2188–2198.

42. Van Wagoner DR, Pond AL, Lamorgese M, Rossie SS, McCarthy PM, Nerbonne JM. Atrial L-type Ca^2+^ currents and human atrial fibrillation. Circ Res. 1999; 85:428–436.

43. Stimers JR, Song L, Rusch NJ, Rhee SW. Overexpression of the large-conductance, Ca^2+^-activated K+ (BK) channel shortens action potential duration in HL-1 cardiomyocytes. PLoS One. 2015;10:e0130588.

44. Rieg AD, Suleiman S, Perez-Bouza A, Braunschweig T, Spillner JW, Schroder T, et al. Milrinone relaxes pulmonary veins in guinea pigs and humans. PLoS One. 2014;9:e87685.

45. Fleming GA, Murray KT, Yu C, Byrne JG, Greelish JP, Petracek MR, et al. M. Milrinone use is associated with postoperative atrial fibrillation after cardiac surgery. Circulation. 2008;118:1619–1625.

46. Bentzen BH, Olesen SP, Renn LC, Grunnet. BK channel activators and their therapeutic perspectives. Front Physiol. 2014;5:389.

