## Supplemental Material for "Conserved Role of the Large Conductance Calcium-Activated Potassium Channel, K_Ca_1.1, in Sinus Node Function and Arrhythmia Risk"

CONTENTS

| | |
|---|---|
| **A. SUPPLEMENTAL METHODS** | |
| Gene sequencing | 3 |
| • PCR | 3 |
| • Sanger sequencing | 3 |
| • Restriction enzyme digestion | 4 |
| Human heart tissue analysis | 4 |
| • Tissue samples | 4 |
| • Immunostaining | 5 |
| • Immunogold electron microscopy | 6 |
| • RNA evaluation | 7 |
| • Western blotting | 8 |
| Murine heart tissue analysis | 9 |
| • Sinoatrial node whole mount immunostaining | 9 |
| • Single cell immunostaining | 10 |
| • Sinoatrial node calcium imaging | 11 |
| Zebrafish studies | 12 |
| • Zebrafish maintenance | 12 |
| • RNA evaluation | 12 |
| • Morpholino injection | 12 |
| • Protein evaluation | 13 |
| • Cardiac function analysis | 13 |
| *Drosophila* studies | 14 |
| • *Drosophila* stocks and crossing | 14 |
| • Lifespan assays | 14 |
| • Nanofluidic qPCR | 15 |
| • Optical heartbeat analysis | 16 |
| • Electrophysiology | 16 |
| Statistical analysis | 17 |
| **B. SUPPLEMENTAL TABLES** | |
| Supplemental Table I | 18 |
| Supplemental Table II | 20 |
| Supplemental Table III | 22 |
| Supplemental Table IV | 23 |
| Supplemental Table V | 24 |
| Supplemental Table VI | 25 |
| Supplemental Table VII | 27 |
| Supplemental Table VIII | 28 |
| **C. SUPPLEMENTAL FIGURE LEGENDS** | 29 |

| | |
|---|---|
| **D. SUPPLEMENTAL FIGURES** | 34 |
| **E. SUPPLEMENTAL REFERENCES** | 48 |

### A. SUPPLEMENTAL METHODS

#### Gene Sequencing

Genomic DNA was obtained from peripheral venous blood samples and protein-coding sequences of the *KCNMA1* gene were PCR-amplified and sequenced.

*PCR.* Intronic primers located at least 40 bp from intron/exon boundaries were designed to amplify 33 exons in the full-length *KCNMA1* transcript (ENST00000286628). The PCR mix contained 100 ng genomic DNA, 200 μM dNTPs (Bioline, Luckenwalde, Germany), 0.3 μM each primer pair, 1.5 mM $MgCl_2$, 0.4-0.8 Unit AmpliTaq Gold® (Applied Biosystems, Waltham, MA) or FastStart Taq (Roche Applied Science, Penzberg, Germany), and 10x PCR reaction buffer. 5x GC-rich buffer (Roche) was added to some reactions to improve amplification. The PCR process was initiated by a denaturation step at 94°C (12 min for AmpliTaq Gold® and 3 min for FastStart Taq), followed by 35 cycles at 94°C (20 sec), 55°C (30 sec), 72°C (60 sec), and a final extension step at 72°C (8 min). Optimal conditions for each primer set were determined (Supplemental Table I) and a negative control was used in every reaction to detect any contamination. PCR products (4 μl) were visualized on 1.5% w/v Agarose (Bioline) in 1x TAE buffer and 50 μg ethidium bromide by gel electrophoresis, then purified using a Montage® $PCR_{96}$ plate (EMD Millipore, Billerica, MA) to remove contaminating salts, unincorporated deoxynucleotides and excess primers.

*Sanger sequencing.* PCR amplicons were subjected to DNA sequence analysis. The sequencing reaction mix (20 μl) contained: 5x dilution buffer (Applied Biosystems), 0.25 μM sequencing primer, 2 μl purified PCR product, and 1 μl BigDye terminator (v3.1, Applied Biosystems). The amplification process was carried out at 96°C (10 sec), 50°C (5 sec), and 60°C (4 min) for 25 cycles after an initial denaturation step at 94°C (5 min). Sequencing primers are listed in Supplemental Table I. The MultiScreen assay system (Millipore) was used to purify each sequencing reaction from contaminating salts, unincorporated primers, and excess dye. Samples were analyzed using the ABI PRISM 3700 DNA Analyser (UNSW Sydney) and sequence electropherograms were analyzed using SeqMan program (DNASTAR Inc, Madison, USA).

*Restriction enzyme digestion*. Restriction enzyme digestion was performed to ascertain the presence or absence of the p.S11_S12delinsG variation identified in Family FF. PCR product (5 µl) was added to a digestion mix (20 µl) containing 5U of MspA1I (New England BioLabs, Ipswich, MA), 10x NEBuffer 4, 2 µl 10x bovine serum albumin (BSA) and incubated at 37°C overnight. Fragments (500-700 bp) were separated by electrophoresis on a 1.5% w/v agarose multi-purpose gel (Bioline) in 1xTAE buffer containing 50 µg ethidium bromide.

**Human Heart Tissue Analysis**

*Tissue samples.* Samples of human right atrial (sinus node, perinodal tissue, atrial myocardium) and right ventricular tissue from unused donor hearts of patients with no history of heart disease were obtained from the Prince Charles Hospital, Chermside, Australia. Right atrial appendage tissue samples were also obtained from patients with non-failing hearts who were undergoing cardiothoracic surgical procedures at St Vincent's Hospital. Patient characteristics are listed in Supplemental Table II. Informed written patient consent was obtained and studies were approved by the Human Research Ethics Committees of the Prince Charles Hospital (EC2565), the University of Manchester, and St Vincent's Hospital (12_164). All work was carried out in accordance with the Human Tissue Act (2004). Immediately after tissue harvesting, samples were fixed and prepared for immunostaining or immunogold electron microscopy, or snap-frozen in liquid nitrogen for subsequent RNA and protein evaluation.

*Immunostaining.* Frozen sinus node tissue sections from 4 human hearts (Supplemental Table II) were selected at approximately 500µm intervals and the precise location of the sinus node and paranodal area were determined using Masson trichrome histology and immunohistochemistry as described.[1] The distribution of $K_{Ca}1.1$ protein within the right ventricle, right atrium, sinus node, and paranodal area was then determined using three rabbit polyclonal IgG antibodies (anti-$K_{Ca}1.1$ APC-021, APC-107, and APC-151, all Alomone Labs, Jersusalem, Israel) targeted to different regions of the protein. Anti-$K_{Ca}1.1$ APC-021 and APC-107 are targeted to intracellular epitopes while anti-$K_{Ca}1.1$ APC-151 is targeted to an extracellular epitope (Supplemental Table III). In order to more accurately determine the precise location of $K_{Ca}1.1$ protein, sections were double-labelled with mouse monoclonal anti-ryanodine receptor 2 (RyR2), anti-caveolin-3, anti-connexin-43, and anti-$Ca_v1.3$ IgG antibodies (Supplemental Table III). The anti-$K_{Ca}1.1$ APC-021 had surface membrane and intracellular staining that co-localized with RyR2 and connexin-43; APC-107 had predominantly intracellular staining that did not co-localize with either RyR2 or connexin 43; APC-151 had surface membrane and intracellular staining that overlapped partially with RyR2 but not with connexin-43 (Supplemental Figure II). Overall, the clearest and most consistent signals were obtained for APC-021 and this antibody was used for subsequent immunostaining and Western blot analyses. Optimal concentrations for the anti-$K_{Ca}1.1$ APC-021 were determined and a negative control was evaluated (Supplemental Figure III).

Tissue sections were fixed in 10% neutral buffered formalin (Sigma-Aldrich, St. Louis, MO) for 30 min and then washed three times (10 min each) in 0.01M phosphate buffered saline (PBS) containing NaCl 0.138 M, KCl 0.027 M, pH 7.4 (Sigma Aldrich). The sections were permeabilized by treatment with 0.1% Triton-X100 (Sigma Aldrich) in PBS for 30 min followed by three PBS washes (10 min each). Sections were blocked using 1% BSA (Sigma Aldrich) in PBS for 60 min, then incubated in primary antibodies diluted in 1% BSA overnight at room temperature. On the following day, the sections were washed three times in PBS (10 min each) and then incubated in cyanine 3 (Cy3)-conjugated donkey anti-mouse IgG and/or fluorescein isothiocyanate (FITC)-conjugated donkey anti-rabbit IgG secondary antibodies (Supplemental Table IV) diluted in 1% BSA for 2 hours. Sections were washed three times in PBS (10 min each) and mounted in Vectashield mounting medium (Vector Laboratories, Burlingame, CA). Sections were imaged using Zeiss LSM5 laser scanning confocal microscope (Carl Zeiss Microscopy, Jena, Germany) using Pascal software (Zeiss Microscopy). The excitation and emission wavelengths used were 490 nm and 520 nm, respectively for FITC (green color), and 552 nm and 565 nm, respectively for Cy3 (red color). Signal intensity measurements were then measured using Volocity software (Improvision, Coventry, England) after background correction.

*Immunogold Electron Microscopy.* Tissues were fixed in 2.5% glutaraldehyde/0.1 M sodium cacodylate buffer, washed, and postfixed with 2% osmium tetroxide and 2% uranyl acetate solution, dehydrated in ethanol series, and embedded in epoxy resin. Sections were viewed by transmission electron microscope (7000, Hitachi, Tokyo, Japan) at a magnification of ×6,000. For immunogold labelling, hearts were perfused with 0.5% glutaraldehyde/2.5% paraformaldehyde in 0.1 M sodium cacodylate buffer (pH 7.2), excised, fixed, and embedded in LR White resin (Polysciences Inc., Warrington, Pennsylvania) as described.[2] Tissue sections were placed on 200 mesh nickel grids and immunolabelled with primary rabbit polyclonal anti-$K_{Ca}$1.1 antibody (APC-021, 1:50), followed by incubation with secondary goat anti-rabbit IgG conjugated with 15-nm gold particles. For double-labelling of $K_{Ca}$1.1 with

RyR2 or $Ca_v1.2$, primary rabbit polyclonal anti-$K_{Ca}1.1$ antibody was used together with mouse monoclonal anti-RyR2 or anti-$Ca_v1.2$ antibody followed by incubation with secondary goat anti-rabbit IgG conjugated with 15 nm gold particles and protein A/G conjugated with 10 nm gold particles, respectively. Negative controls for the APC-021 labelling are shown in Supplemental Figure III. Immunogold-labelled sections were stained with uranyl acetate and lead citrate and examined by a JEM-1400 transmission electron microscope (Jeol, Peabody, Massachusetts).

*RNA evaluation.* $K_{Ca}1.1$ isoforms were evaluated in atrial tissue using quantitative RT-PCR (qPCR) analysis. Total RNA was isolated from frozen heart tissues using RNeasy Micro kit (Qiagen, Venlo, Netherlands) according to the manufacturer's protocol. RNA concentration was measured using a Nanodrop spectrophotometer (Labtech International, Uckfield, UK). 1 ug of total RNA from each sample was reverse-transcribed with Superscript III Reverse Transcriptase Supermix (Invitrogen, Carlsbad, CA) in a 20 μl reaction according to the manufacturer's instructions using random hexamer priming. Aliquots of the resulting cDNA were diluted 20-fold in water for direct use in qPCR.

The relative abundance of selected cDNA fragments was determined with qPCR using a Light-Cycler instrument (Applied Biosystems) with SYBR-Green Master Mix (Roche) and forward and reverse primers (0.6 μM each) under the following conditions: denaturation at 95°C (3 min); 35 cycles at 94°C (1 min), at 60°C (1 min) and at 72°C (1 min). Hypoxanthine phosphoribosyltransferase 1 (Hprt) and β-actin were used as an internal reference in each reaction. Amplification was followed by melting curve analysis using the program run at the step acquisition mode to verify the presence of a single amplification product in DNA. Accumulation of PCR products and the threshold cycle were monitored and determined using the LightCycler analysis software. For all mRNAs in each sample, at least three separate measurements were made with 1 μl aliquots of each cDNA sample.

Oligonucleotide primers for qPCR analysis for each transcript were:

| | |
|---|---|
| Insertless (F) | CCATTAAGTCGGGCTGATTTAAG |
| Insertless (R) | CCTTGGGAATTAGCCTGCAAGA |
| STREX (F) | GCCAAGATGTCCATCTACAAG |
| STREX (R) | GCACGGAAACTGGTGGAGCAA |
| SV27 (F) | CTAAGCCGGGCAAGTTG |
| SV27 (R) | GAGTCCAGGACACTGACG |
| DEC (F) | GGTTTACAGATGAGCCGGATA |
| DEC (R) | CATCTTCAACTTCTCTGATTGG |
| Del e23 (F) | GACGTCACAGATCCCAAAAGAAT |
| Del e23 (R) | TGATCATTGCCAGGAATTAACAA |
| Total *KCNMA1* (F) | CCATTAAGTCGGGCTGATTTAAG |
| Total *KCNMA1* (R) | CCTTGGGAATTAGCCTGCAAGA |
| β-actin (F) | TCCTTCGTTGCCGGTCCACA |
| β-actin (R) | CCTCTCTTGCTCTGGGCCTCG |
| Hprt (F) | TGAGGATTTGGAAAGGGTGT |
| Hprt (R) | TAATCCAGCAGGTCAGCAAA |

*Western blotting.* For extraction of total protein from frozen heart tissues, samples were crushed under liquid nitrogen and re-suspended in homogenization buffer (10 mM EDTA, 300 mM sucrose, 0.35 mM SDS and protease cocktail inhibitors; Sigma-Aldrich). Samples were spun at 4°C for 6 min at 10,000 rpm. Supernatant was removed and re-suspended in an equal volume of Laemmli buffer (250 mM Tris-HCl pH 6.8, 5% SDS, 40% glycerol, 5% ß-mercaptoethanol and 0.005% bromophenol blue) for protein gel analysis. A 10 μl aliquot of sample was removed for estimation of protein concentration using a Bradford colorimetric assay (BioRad Laboratories, Hercules, CA). The samples were stored in aliquots at -80°C. Prior to loading on the gel, samples were heated to 80°C for 5 min then 20 μl proteins from each sample were loaded. Proteins were separated by SDS-PAGE using 10% gels (125 mM Tris-HCL, 960 mM glycine, 0.5% (w/v) SDS, pH 8.3). The gels were run at 90 V for 2-3 h. Gels were transferred to a PVDF (GE Healthcare, Little Chalfont, UK) membrane, at 30 V overnight at 4°C in transfer buffer (125 mM Tris-HCL, 960 mM glycine, 0.5% (w/v) SDS, pH 8.3). The blots were blocked (3% BSA, Tween 20 in TBS) for 1 h and incubated with rabbit polyclonal anti-$K_{Ca}$1.1 antibody (APC-021, 1:200) either overnight at 4°C or for 2 h at room temperature. A negative control experiment in which the procedure was undertaken without the primary APC-021 antibody was also performed (Supplemental Figure IV). Blots were washed with TBS-Tween 20, incubated in horseradish peroxidase (HRP)-conjugated goat anti-rabbit IgG secondary antibody for 1 h and then exposed to chemiluminescent substrate (Pierce Biotechnology, Rockford, IL). Blots were normalized to β-tubulin and hybridization signals were quantified with ImageJ software.

**Murine Heart Tissue Analysis**

*Sinoatrial node whole mount immunostaining*. The sinoatrial node (SAN) region was dissected from the right atrium of adult wild-type mice and fixed in 4% paraformaldehyde for 30 min at 4°C. Whole-mount immunostaining was performed using rabbit polyclonal anti-$K_{Ca}$1.1 (APC-021), mouse monoclonal anti-ryanodine receptor 2 (RyR2), mouse monoclonal anti-connexin-43, and mouse monoclonal anti- $Ca_v$1.3 IgG antibodies (Supplemental Table III). Dissected SAN complexes were permeabilized by washing for 30 min in PBS containing 1% Triton X-100. After blocking of non-specific binding by incubation for 1 h with 0.5% Triton X-100 in PBS containing 10% normal goat serum (NGS), SAN whole-mount preparations were incubated at 3 days at 4°C with the primary antibodies (Supplemental Table III). They were then washed three times with PBS containing 0.1% Triton X-100, incubated for 4 h with Cy3-conjugated goat antibodies to rabbit IgG (Supplemental Table IV), and washed three times with PBS containing 0.1% Triton X-100. All antibodies were diluted with 0.5% Triton X-100 in PBS containing 5% NGS. Whole-mounts were mounted in ProLong Gold antifade reagent with DAPI (Invitrogen), a water-soluble nuclear and chromosomal counterstain. The SAN preparations were mounted with the endocardium uppermost. Immunofluorescence was examined with a confocal microscope (Leica TCS NT). DAPI, FITC, and Cy3 fluorescence was excited at 364, 488 and 543 nm, respectively. Images were processed with Image J software. For controls, pre-incubation of the primary antibodies with their peptide antigen, or incubation of the SAN whole mounts with only the secondary antibody, reduced fluorescence to non-detectable levels. The lack of detectable fluorescence confirmed the specificity of the antibody binding.

*Single cell immunostaining*. Single pacemaker cells were enzymatically isolated from the SAN region of adult mouse hearts. The SANs were dissected free and placed into calcium-free Tyrode solution containing (mM): NaCl 133, KCl 4.0, $NaH_2PO_4$ 1.2, $MgCl_2$ 1.2, glucose 10, HEPES (N-2-hydroxyethylpiperazine-N'-2-ethansulphonic acid) 10, for 5 min. Intact SAN tissue was then transferred into 25 μmol/L calcium-Tyrode solution containing 1 mg/ml collagenase (Worthington Type II) at 35°C for 60 min before it was transferred into fresh 25 μmol/L calcium-Tyrode solution containing 0.5 mg/ml elastase (Sigma-Aldrich) for a further 50 min. The SAN was then placed in KB solution containing (mM): KCl 50, KATP 5, glucose 10, taurine 20, creatine 5, glutamic acid 5, succinic acid 5, HEPES 5, $KH_2PO_4$ 20, $MgSO_2$ 5, pyruvic acid 5, K-EGTA 0.04; pH was adjusted to 7.2 with KOH. SANs were kept in KB solution for at least 4 h at 4°C, before being triturated gently to dissociate the cells. The cells were then placed onto glass coverslips and maintained at room temperature for about 1 h in order to allow cells to adhere to the glass. Isolated single pacemaker cells were fixed in 4% paraformaldehyde for 1 min then permeabilized by incubating for 5 min in PBS containing

NGS, 1% BSA, and 0.1% Triton X-100. After blocking of non-specific binding sites by incubation for 30 min with 0.01% BSA in PBS containing 10% NGS, single pacemaker cells were exposed to primary antibodies (Supplemental Table III).

*Sinoatrial node $Ca^{2+}$ imaging.* Mice were sacrificed using $CO_2$ chambers. The hearts along with lungs and thymus were dissected and washed with Tyrode solution (140mM NaCl, 5.4mM KCl, 5mM Hepes, 5.5mM Glucose, 1mM $MgCl_2$, 1.8mM $CaCl_2$, pH7.4, oxygenated and warmed to 37°C). The tissue was transferred to a silicon-coated dish containing warmed Tyrode solution. The right atrium (RA) along with inferior vena cava (IVC), superior vena cava (SVC) and part of the interatrial septum were dissected from the rest of the tissues. Once completely separated, the tissue was re-pinned through the IVC, SVC and RA and any remaining ventricular tissue was removed. The anterior walls of the RA, IVC and SVC were opened using dissection scissors. The pins were repositioned to stretch the tissue gently revealing the SAN region, which lies between the interatrial septum, RA, IVC and SVC. The dissected SAN was transferred to a silicon-coated petri dish with ~2ml of warmed Tyrode solution. Tissue was pinned to the new dish with the endocardial surface facing up. To stain the tissue, 10µM Cal-520 dye (AAT Bioquest, Sunnyvale, CA) in 2ml Tyrode solution was added and the tissue was incubated for 1 h shaking gently at room temperature in the dark. Afterwards, the tissue was washed three times with dye-free warmed Tyrode and then incubated for 30 min in 2 ml of warmed Tyrode containing 5µM blebbistatin (Sigma-Aldrich). The tissue was turned around so that the endocardial surface was facing down. Calcium imaging was performed on the Eclipse Ti2 inverted microscope (Nikon Corp., Tokyo, Japan) while perfused at a rate of 12 ml/min with warmed Tyrode containing 5µM blebbistatin. The Tyrode solution was kept at 37°C throughout the imaging process. $Ca^{2+}$ transient beating rate was assessed at baseline, and at 7 time points of 5s intervals after addition of the $K_{Ca}1.1$ antagonist, paxilline (10 μM; Sigma-Aldrich), ± isoproterenol (500 nM; Sigma-Aldrich). The SAN preparation was subjected to 5 min wash periods with warmed Tyrode's solution between each drug administration.

**Zebrafish Studies**

*Zebrafish maintenance.* Wild-type TE zebrafish were maintained under standard aquarium conditions and in compliance with the Australian Code of Practice for the Care and Use of Animals for Scientific Purposes. Procedures involving zebrafish were carried out in accordance with institutional biosafety regulations and with approval from the Garvan Institute of Medical Research/St Vincent's Hospital Animal Ethics Committee.

*RNA evaluation.* Hearts from 3-day post fertilization (dpf) zebrafish embryos were isolated as previously described.[3] RNA from pooled embryonic hearts or whole embryos was then purified using Qiazol reagent (Qiagen, Chadstone, Australia) and cDNA was prepared using a SuperScriptIII kit according to the manufacturer's instructions (Invitrogen, Mount Waverley, Australia). For detection of *kcnma1a* and *kcnma1b* expression, cDNA was amplified by reverse transcription-polymerase chain reaction (RT-PCR) using the gene-specific primers listed in Supplemental Table V.

*Morpholino injections.* Morpholino antisense oligonucleotides were designed targeting the translation start sites of zebrafish *kcnma1a* and *kcnma1b* (Gene Tools, Oregon, USA). These morpholinos or a standard Gene Tools control morpholino, were dissolved in water to a concentration of 150μM, and a volume of 2nL was injected into embryos at the 1-2 cell-stage using a Picospritzer III pressure injector (Parker Hannifin, Castle Hill, Australia). Morpholino sequences are shown in Supplemental Table V.

*Protein evaluation*. To assess the efficacy of inhibition of protein translation by the *kcnma1* morpholino, total protein extracts from morpholino-injected embryos were separated using SDS polyacrylamide gel electrophoresis and Western blotting was carried out as described above for human tissues, using two primary rabbit polyclonal antibodies: anti-$K_{Ca}$1.1 antibody specific for the *kcnma1b* zebrafish ortholog (APC-021, 1:300) and an anti-$K_{Ca}$1.1 antibody that binds both *kcnma1a* and *kcnma1b* isoforms (AB9770, 1:400), with HRP-conjugated goat anti-rabbit IgG secondary antibody (1:2000).

*Cardiac function analysis***.** Heart rate, maximal atrial diameter, atrial fractional area change (FAC) and ventricular fractional shortening (FS) were determined at 3 dpf.[3] Embryos were lightly anesthetized in 50 mg/L tricaine at room temperature. Heart rate was measured by direct microscopic observation. Each embryo was observed twice, for a period of 15 sec each time, separated by an interval of 15 min. The mean of the two counts was used to calculate heart rate. Atrial area and ventricular diameter were measured by video analysis using a Nikon DS-Qi1MC camera (Nikon) with NIS-Elements AR v3.1 software mounted on a Leica DM-IL inverted microscope (Leica Microsystems, North Ryde, Australia). Embryos were mounted side-on on a depression slide in 3% methylcellulose, short videos were recorded and measurements were taken over 6 consecutive cardiac cycles. Atrial maximal diameter was measured using still images at end-diastole. For measuring atrial FAC, the atrium was traced and the area calculated using the free drawing function of the software, on still images of end-systole and end-diastole. The formula %FAC = (diastolic area − systolic area) / (diastolic area) × 100 was used. Similarly, ventricular FS was calculated by measuring the diameter of the short axis at end-systole and end-diastole, then using the formula %FS = (diastolic diameter − systolic diameter) / (diastolic diameter) × 100.[4] For all parameters, the mean of measurements over 6 consecutive cycles was used. Results are presented as mean ± SEM.

Values were compared using Student's t-test, with a value of $p<0.05$ considered statistically significant.

**Drosophila Studies**

*Drosophila Stocks and Crossing*. The fly lines *slo*$^{4}$, *slo*$^{1}$, *Df (3R) BSC 397, w*$^{1118}$*, pan-neural driver ElAV-Gal4,* and *w; Cyo/Sco; TM3/TM6B* were obtained from the *Drosophila* Stock Center. The *slowpoke* UAS-RNAi line (construct ID: 108671) and control RNAi insertion *y,w[1118];P{attP,y[+],w[3`]}* were from the Vienna *Drosophila* RNAi Center. The cardiac tissue-specific Hand4.2-Gal4 driver was a gift from Zhe Han and Eric Olson.[5] UAS-KCNMA1 flies were generated from cloned human cDNA that was cloned into pcDNA3.1, then the pUAST vector, and finally PhiC31-mediated transformation via injection in *w*$^{1118}$ strain. PBac *slo* genomic rescue flies were generated using a bacterial artificial chromosome construct containing the *slo* locus (PBac slo, gift of Hugo Bellen)[6] which was injected into w$^{1118}$ flies, transformed via PhiC31, then crossed to slo$^{4}$/Df(3R) BSC 397 flies. The cardiac-specific inducible HandGS-Gal4 driver[7,8] was a gift from Laurent Perrin. Flies were collected two days from eclosion and maintained on transgene-inducing food containing mifipristipne (RU486) from 2 days until 3 weeks of age. Flies were reared on a standard yeast/molasses/cornmeal diet and maintained at 25°C. Flies were collected within 24 h of eclosion and placed into vials with 20-25 same sex cohorts. Vials were changed every 2-3 days.

*Lifespan assays*. Female progeny were collected for 3 days. Then, they were briefly anesthetized and separated in groups of 25 flies in each vial. The flies were kept at 25°C, and the dead flies were counted every 2 days after transfer. The experiment was performed on 200 flies. Data were analyzed using Prism 5.0 (GraphPad Software, La Jolla, CA).

*Nanofluidic qPCR.* Five fly hearts were isolated and snap frozen in 10μl of water. Each sample was lyophilized and brought up to a final volume of 40μl of lysis buffer (0.25% NP40 in water). Lysis was carried out by heating individual hearts in lysis buffer for 2 min at 98°C. The lysed hearts were then briefly centrifuged, and 30μl transferred to a fresh tube. Reverse transcription was carried out by using 3.3μl of lysed heart (~8% of a single heart) in a 5μl final reaction volume using the VILO Reaction mix as per the manufacturer's instructions (Life Technologies, Carlsbad, CA), resulting in a final volume of 6μl of RT cDNA per heart. For use with the BioMark platform, we then carried out pre-amplification for 10 genes using 3μl Master Mix (Fluidigm Corp., San Francisco, CA), 1.5μl 10x single target amplification, 100μM each primer (listed below), 0.5 M EDTA (pH 8, 0.075μl), and water, to a final volume of 9μl. Pre amplification was then carried out in a volume of 15 μl (pre-amp cycling 95°C for 2 min, and 20 cycles of 96°C 5 sec, 60°C for 4 min). For removal of single stranded DNA prior to nanofluidic cycling, 6 μl of Exosap solution (4.2μl water, 0.6μl Exonuclease 1 Rn Buffer, Exonuclease 1 [20 units/μl, New England Biolabs]) was added to the 15μl final reaction volume of the RT step. The resulting 21 μl final volume was then incubated at 37°C for 30 min, and then heat inactivated at 80°C for 15 mins. The pre-amplified volume (21μl) was then diluted 10-fold in DNA suspension buffer (Teknova Inc., Hollister, CA), and stored at -20°C prior to running on chip. Pre-amplified products for each individual fly heart were then assayed using 48.48 nanofluidic qPCR arrays on a Biomark system (Fluidigm). EvaGreen DNA binding dye (Biotium, Hayward, CA) was used to detect amplified product.

Primers used for pre-amplification:

Ca-α1D- F:CTACGTCCACTGCGACTTGTA; R:AGTGGCACCATGGCCTTTAA

Elk- F:CTGCCCTTTGATCACCTGTAC; R:CAGGAGACGCGTCAATTTCA

Ih- F: ACAACCGACTGGCCATGA; R:GTGCCCGGAAGTTTTCTGAC

Irk1- F: GCAACGTTGTGCAGGGAAA; R:CGTCAACCAGGGTGGTGAA

KCNQ- F:TGAAGCCCTACGACGTCAA; R:GCATTTTAACGCGACCCAAC

Seizure- F:AATCCAGAGAGCCGGCAATA; R:CCGACCGTTGGGTAAATACAC

Shaker- F:CCGAGCTTCGATGCGATTTTA; R: GGGACATTGACCGGTCTCC

Slowpoke- F:TCATCCAGCTGATGCAGTACC; R:ATCGTCGCCCTGTTTCCAA

SUR- F:GCAGCTGAAGGAGTTTGTCA; R:AGGTTTAGCCCTCCATCACA

*Optical Heartbeat Analysis*. Adult heart parameters were analyzed as described previously.[4,9] Briefly, flies were dissected in an artificial hemolymph containing 108 mM $Na^+$, 5 mM $K^+$, 2 mM $Ca^{2+}$, 8 mM $MgCl_2$, 1 mM $NaH_2PO_4$, 4 mM $NaHCO_3$, 10 mM sucrose, 5 mM trehalose, and 5 mM Hepes (pH 7.1). All reagents were obtained from Sigma-Alrich. Movies of beating hearts were recorded using a Leica S9300 microscope with a water immersion lens, high-speed digital camera (Hamamatsu EM-CCD) using HC Image software (Hamamatsu Photonics Corp., Hamamatsu, Japan). Heart function parameters including heart period, heart diameters, systolic and diastolic intervals, FS and arrhythmia index were quantified using our semi-automated optical heartbeat analysis software.[4]

*Electrophysiology.* Semi-intact heart preparations were incubated in artificial hemolymph containing 10 µM blebbistatin (Sigma-Aldrich) and equilibrated with oxygenation in the dark for 45-60 min until the hearts stopped beating. The preparation was then supplied with fresh saline without blebbistatin and electrical potentials were recorded from the conical chamber using sharp glass electrodes (20-50 MΩ) filled with 3M KCl and standard electrophysiological techniques. Data were acquired using an Axon-700B multiclamp amplifier, signals were digitized using the DIGIDATA 1322A and data were captured and analyzed using PClamp 9.0 and Clampfit 10.0 software respectively (all from Molecular

Devices, Sunnyvale, CA). Data was quantified from representative 30 s recordings where the resting membrane potential had remained stable for at least 30 s.

**Statistical Analysis**

Differences between groups were assessed using Student's t test or ANOVA. Data are expressed as mean ± SEM. A p value <0.05 was considered statistically significant.

#### B. SUPPLEMENTAL TABLES

**Supplemental Table I.** Primers for *KCNMA1* Screening.

| Exon | Primer ID | Primer sequence | PCR size | PCR condition | Sequencing Primer |
|---|---|---|---|---|---|
| 1 | KMA1x1F | gcggctcgctgtatatatctc | 836 | Fast Start Taq, GC buffer | KMA1x1F |
| | KMA1x1R | aaacaagaggacaggattgagc | | | KMA1x1R |
| 2 | KMA1x2F | cccttctggctctggttctt | 335 | Amplitaq Gold | KMA1x2F |
| | KMA1x2R | cctggggaatatcacgtctc | | | |
| 3 | KMA1x3F | ggatgagggtgagatttcca | 292 | Amplitaq Gold | KMA1x3R |
| | KMA1x3R | gcaaaggttggtgtcaaggt | | | |
| 4 | KMA1x4F2 | ggaggcagaaagaccaaatg | 330 | Amplitaq Gold | KMA1x4R2 |
| | KMA1x4R2 | tcttgactgcgagagcagag | | | |
| 5 | KMA1x5F | tgtacattggatgccagcac | 244 | Amplitaq Gold | KMA1x5R |
| | KMA1x5R | tgcaaattctgtccttcagc | | | |
| 6 | KMA1x6F | cgacccttgggaactgtaaa | 246 | Amplitaq Gold | KMA1x6R |
| | KMA1x6R | atgaaagcaagcacacctga | | | |
| 7 | KMA1x7F | ttagatgtggcagcctctcc | 317 | Amplitaq Gold | KMA1x7R |
| | KMA1x7R | ttgcttcccatcttcctgtc | | | |
| 8 | KMA1x8F | ctcgtgtggagagtggtgtg | 448 | Amplitaq Gold | KMA1x8F |
| | KMA1x8R | tcaccatcgtgtttggattt | | | |
| 9 | KMA1x9F | ccttttctttggcctgtcttt | 250 | Amplitaq Gold | KMA1x9F |
| | KMA1x9R | tgcctacatgcatgaaacaaa | | | |
| 10 | KMA1x10F | gatgggagctgggcttttat | 396 | Amplitaq Gold | KMA1x10F |
| | KMA1x10R | ccaaaagggatcatggctta | | | |
| 11 | KMA1x11F | gcaagcaaaagggtgactgt | 322 | Amplitaq Gold | KMA1x11R |
| | KMA1x11R | gtgtcccctcagcacagagt | | | |
| 12 | KMA1x12F | ttccccaagaagctgtcaac | 280 | Amplitaq Gold | KMA1x12R |
| | KMA1x12R | gggcaaaaaggcctcataga | | | |
| 13 | KMA1x13F | gggcttgtccagtgaaaagt | 326 | Amplitaq Gold | KMA1x13F |
| | KMA1x13R | acctgtggatgggtcttcag | | | |
| 14 | KMA1x14F | aagggaaaagaagggcaaga | 377 | Amplitaq Gold | KMA1x14F |
| | KMA1x14R | cccgtcgatctgttttgagt | | | |

| | | | | | |
|---|---|---|---|---|---|
| 15 | KMA1x15F | ctcaggtttcccttttacagc | 235 | Amplitaq Gold | KMA1x15R |
| | KMA1x15R | gcaaggggcacattcaata | | | |
| 16 | KMA1x16F | ggatcccctcggtaatgact | 237 | Amplitaq Gold | KMA1x16F |
| | KMA1x16R | gaacgcactctcaccatcaa | | | |
| 17 | KMA1x17F | gacaaaatctttggggaaagc | 290 | Amplitaq Gold | KMA1x17R |
| | KMA1x17R | gcctacttccgtgggtcaa | | | |
| 18 | KMA1x18F | accgtggaggaaatgtggta | 222 | Amplitaq Gold | KMA1x18R |
| | KMA1x18R | gggaaggaaaaggaatttgg | | | |
| 19 | KMA1x19F | ccttgctgtgtgtgaccttg | 313 | Amplitaq Gold | KMA1x19R |
| | KMA1x19R | acagcacaagacagggatga | | | |
| 21 | KMA1x21F | tgagccttgagtgtgtgtcc | 343 | Amplitaq Gold | KMA1x21R |
| | KMA1x21R | ccagcctttaagaagccatc | | | |
| 22 | KMA1x22F | aatggccttcagtgacaacc | 330 | Amplitaq Gold | KMA1x22F |
| | KMA1x22R | ccaactagggaaaccccatt | | | |
| 23 | KMA1x23F | gcacatagtaagctctcagcaaa | 447 | Amplitaq Gold | KMA1x23F |
| | KMA1x23R | gaagcacaggcatgatcaaa | | | |
| 24 | KMA1x24F | ccctctcctctcacttttgc | 384 | Amplitaq Gold | KMA1x24R |
| | KMA1x24R | caggctgatgtgagcctctt | | | |
| 25/26 | KMA1x25F | cccgggaatactaattgttga | 1185 | Amplitaq Gold | KMA1x25F |
| | KMA1x26R | caccaacaacagaacaaaagga | | | KMA1x26R |
| 27 | KMA1x27F | ttccatgcttttgtttccttc | 324 | Amplitaq Gold | KMA1x27F |
| | KMA1x27R | tcaaaggttggaggtgctct | | | |
| 28 | KMA1x28F | cggctgtcatgactttaggg | 363 | Amplitaq Gold | KMA1x28F |
| | KMA1x28R | ggagaaaagccagatgccta | | | |
| 29 | KMA1x29F | tgatgaaaagacccgtttcc | 364 | Amplitaq Gold | KMA1x29F |
| | KMA1x29R | ttcccatcacctccaaactc | | | |
| 30 | KMA1x30F | ggctggcctcctttattgag | 504 | Amplitaq Gold | KMA1x30F |
| | KMA1x30R2 | ggggaaatgagtggcagata | | | |
| 31 | KMA1x31F | tgccagtaaagtgctcaaca | 397 | Amplitaq Gold | KMA1x31F |
| | KMA1x31R | tagcgatagcagcagcacag | | | |
| 32 | KMA1x32F | ctcactgtcggcttctatcg | 689 | Fast Start Taq, GC buffer, | KMA1x32F |
| | KMA1x32R | ggtggtgaccatcattctcc | | annealing temp: 61.3$^{o}$C | |

**Supplemental Table II.** Clinical details of heart tissue donors.

| Patient | Age | Sex | Indication for surgery | Heart disease | AF history | Tissue obtained |
|---|---|---|---|---|---|---|
| 1 | 31 | M | Heart Tx donor: CH, CR arrest | Nil known | No | Left ventricle |
| 2 | 40 | F | Heart Tx donor: CH, BD | Nil known | No | Sinus node + adjacent atrial muscle |
| 3 | 40 | F | Heart Tx donor: CH, BD | Nil known | No | Sinus node + adjacent atrial muscle |
| 4 | NA | NA | Heart Tx donor | Nil known | No | Sinus node + adjacent atrial muscle |
| 5 | NA | NA | Heart Tx donor | Nil known | No | Sinus node + adjacent atrial muscle |
| 8 | 42 | M | AVR | IE | No | Right atrial appendage |
| 24 | 50 | M | CABG | CAD | No | Right atrial appendage |
| 43 | 57 | M | CABG | CAD | No | Right atrial appendage |
| 22 | 58 | M | CABG | CAD | No | Right atrial appendage |
| 29 | 59 | M | CAG, AVR | CAD, AS, AR | No | Right atrial appendage |
| 13 | 62 | M | CABG | CAD | No | Right atrial appendage |
| 18 | 63 | M | Bentall's procedure | AAA | No | Right atrial appendage |
| 46 | 63 | M | CABG | CAD | No | Right atrial appendage |

| | | | | | | |
|---|---|---|---|---|---|---|
| 31 | 73 | F | CABG | CAD | No | Right atrial appendage |
| 51 | 78 | M | CABG | CAD | No | Right atrial appendage |
| 42 | 81 | F | CABG | CAD | No | Right atrial appendage |
| 60 | 60 | M | CABG | CAD | Yes | Right atrial appendage |
| 7 | 61 | F | CABG, AVR | CAD, AS, AR | Yes | Right atrial appendage |
| 33 | 61 | M | CABG | CAD | Yes | Right atrial appendage |
| 28 | 73 | F | CABG, AVR | CAD, AS | Yes | Right atrial appendage |
| 34 | 76 | F | CABG, MVR, TVR, LA reduction | CAD, MR, TR | Yes | Right atrial appendage |
| 14 | 81 | M | AVR | AS | Yes | Right atrial appendage |
| 17 | 81 | M | CABG, AVR | CAD, AS | Yes | Right atrial appendage |
| 41 | 81 | F | CABG | CAD | Yes | Right atrial appendage |

AAA, ascending aortic aneurysm; AF, atrial fibrillation; AR, aortic valve regurgitation; AS, aortic valve stenosis; AVR, aortic valve replacement; BD, brain death; CABG, coronary artery bypass grafts; CAD, coronary artery disease; CH, cerebral haemorrhage; CR, cardiorespiratory arrest; IE, infective endocarditis; LA, left atrium; MR, mitral valve regurgitation; MVR, mitral valve replacement; NA, not available; TR, tricuspid valve regurgitation; TVR, tricuspid valve replacement; Tx, transplant.

**Supplemental Table III.** List of primary antibodies.

| Antigen | Host species | Ab type | Isotype | Dilution | Source/catalogue no. |
|---|---|---|---|---|---|
| Caveolin-3 | Mouse | Monoclonal | IgG | 1:500 | BD Biosciences, 610421 |
| L-type $Ca^{2+}$ channel, $Ca_v1.2$ | Mouse | Monoclonal | IgG | 1:50 | Alomone Labs, ACC-003 |
| L-type $Ca^{2+}$ channel, $Ca_v1.3$ | Mouse | Monoclonal | IgG | 1:100 | Abcam, ab85491 |
| Connexin-43 | Mouse | Monoclonal | IgG | 1:50 | Millipore, MAB3068 |
| $K_{Ca}1.1$ (extracellular, 199-213) | Rabbit | Polyclonal | IgG | 1:50 to 1:800 | Alomone Labs, APC-151 |
| $K_{Ca}1.1$ (intracellular, 1097-1196) | Rabbit | Polyclonal | IgG | 1:50 to 1:800 | Alomone Labs, APC-021 |
| $K_{Ca}1.1$ (intracellular, 1184-1200) | Rabbit | Polyclonal | IgG | 1:50 to 1:800 | Alomone Labs, APC-107 |
| $K_{Ca}1.1$ (intracellular, 1184-1200) | Rabbit | Polyclonal | IgG | 1:500 | Millipore, AB9770 |
| Ryanodine receptor-2 | Mouse | Monoclonal | IgG | 1:100 | Thermo Scientific, MA3-916 |

**Supplemental Table IV.** List of secondary antibodies and other reagents for indirect immunolabelling.

| Host Species | Type | Isotype | Conjugate | Dilution | Source |
|---|---|---|---|---|---|
| Goat | Mouse | IgG | Cy3 | 1:400 | Millipore, AP181C |
| Donkey | Mouse | IgG | Cy3 | 1:400 | Millipore, AP192C |
| Donkey | Guinea pig | IgG | Cy3 | 1:400 | Millipore, AP193C |
| Goat | Rabbit | IgG | FITC | 1:100 | Millipore, 12-507 |
| Donkey | Rabbit | IgG | FITC | 1:100 | Millipore, AP182F |
| Goat | Rabbit | IgG | HRP | 1:3000 | GE Healthcare, NA934V |
| Goat | Rabbit | IgG | 15 nm gold | 1:50 | SPI Supplies, 4538-AB |
| Protein A/G | | | 10 nm gold | 1:50 | SPI Supplies, 4233-AB |

**Supplemental Table V.** Primers used for RT-PCR and morpholino oligonucleotides used for *kcnma1a* and *kcnma1b* down-regulation in zebrafish embryos.

| | |
|---|---|
| *Primers* | |
| *kcnma1a*_F | TAATACGACTCACTATAGGGAAAAATCCGGACTCTTCTGTCTC |
| *kcnma1a*_R | ATTTAGGTGACACTATAGATTGTGCCATCAGCCTGAATA |
| *kcnma1b*_ F | TAATACGACTCACTATAGGGACGTGCATCGCGTCAGTA |
| *kcnma1b*_R | ATTTAGGTGACACTATAGATTGTCGAAATGCTCACAGGA |
| *actb1*_F | CAGACATCAGGGAGTGATGGTT |
| *actb1*_R | CTGTGTCATCTTCTCTCTGTTGG |
| *Morpholino oligonucleotides* | |
| *kcnma1a* ATG | CTAGAGCTGCTGCTGCCATAGGATC |
| *kcnma1b* ATG | GTTATCAGATCAGATTCAGTCATCT |
| Control | CCTCTTACCTCAGTTACAATTTATA |

**Supplemental Table VI.** *KCNMA1* sequence variants identified in 118 probands with familial AF.

| GRCh38/ hg38 genomic start position | Transcript effect | Protein effect | Minor allele frequency | | dbSNP | ClinVar |
|---|---|---|---|---|---|---|
| | | | AF cohort | gnomAD (All/NFE) | | |
| 77637744 | c.-102_-101insGGCAGC | | 0.182 | Absent* | Absent | Not listed |
| 77637619 | c.24C>T | p.Gly8Gly | 0.004 | 0.00012/0.00022 | rs748427000 | LB/VUS |
| 77637612 | c.31_36delAGCAGCinsGGC | p.Ser11_Ser12del;insGly | 0.004 | Absent | Absent | Not listed |
| 77637528 | c.114_115insTCC | p.Ala38_Ser39insSer | 0.008 | Absent | Absent | Not listed |
| 77404046 | c.379-23delAinsTTTT | | 0.004 | Absent | Absent | Not listed |
| 77185039 | c.603-123delG | | 0.008 | Absent* | Absent | Not listed |
| 77185005 | c.603-89C>T | | 0.008 | Absent* | Absent | Not listed |
| 77184832 | c.687C>T | p.Phe229Phe | 0.297 | 0.4328/0.3509 | rs1131824 | B/LB |
| 77112523 | c.885-81G>A | | 0.008 | Absent* | rs12778504 | Not listed |
| 77112319 | c.960+48C>A | | 0.008 | 0.02239/0.02758 | rs12355548 | Not listed |
| 77108446 | c.1223+35C>T | | 0.068 | 0.05844/0.09194 | rs45459294 | Not listed |
| 77090562 | c.1224-52G>C | | 0.008 | 0.02224/0.02765 | rs11002011 | Not listed |

| | | | | | | |
|---|---|---|---|---|---|---|
| 77079626 | c.1524-76G>T | | 0.068 | 0.05819/0.09068 | rs45495698 | Not listed |
| 77073312 | c.1594-60A>G | | 0.055 | 0.05845/0.09077 | rs2274416 | Not listed |
| 77018976 | c.2015+37C>T | | 0.699 | 0.5510/0.6782 | rs16934182 | Not listed |
| 77012074 | c.2016-31A>G | | 0.004 | 0.00058/0.00096 | rs201129775 | Not listed |
| 76969911 | c.2484+63C>T | | 0.792 | 0.8187/0.7744 | rs2288837 | Not listed |
| 76949407 | c.2485-41C>T | | 0.004 | 0.01029/0.009107 | rs79866794 | Not listed |
| 76949325 | c.2526C>T | p.Val842Val | 0.013 | 0.009098/0.01395 | rs41274568 | B/LB |
| 76914863 | c.3016+73G>T | | 0.008 | 0.009839/0.01159 | rs2288838 | Not listed |
| 76909941 | c.3147+25T>A | | 0.008 | 0.01022/0.004786 | rs141315319 | Not listed |
| 76891672 | c.3195C>T | p.Thr1065Thr | 0.013 | 0.00576/0.00940 | rs45527834 | B/LB |
| 76889586 | c.3343-17C>A | | 0.004 | 0.00007/0.00015 | rs199571730 | Not listed |
| 76889362 | c.3461+89T>A | | 0.004 | Absent* | Absent | Not listed |

Numbering is based on *KCNMA1* transcript ENST00000286628, NM_001161352.2. B, Benign; gnomAD, Genome Aggregation Database (v3, accessed June 2020). LB, Likely benign; NFE, non-Finnish Europeans; VUS, variant of uncertain significance.
*Low coverage of this region.

**Supplemental Table VII**. Clinical characteristics of *KCNMA1* mutation carriers in Family FF.

| ID/sex | Age (yr) at study/ AF dx | AF | ECG | LAD (mm) | LVEF (%) | Additional information |
|---|---|---|---|---|---|---|
| I-1/M | 84 | Suspected | SR (65 bpm), APC, LAHB, RBBB | NA | NA | Palpitations, no documented AF; Holter: SR, rate 45-132 bpm; bifascicular block. No cardiac medications. |
| II-1/M | 56/50 | Permanent | AF | 47 | 40 | Tachy-brady syndrome, PPM; Holter: AF, rate <30 to >200 bpm, pauses (up to 2.9 s); stress test: early tachycardic response to exercise (max 190 bpm). On warfarin, metoprolol, flecainide; adequate rate control difficult to achieve. |
| II-3/M | 49/34 | Permanent | AF | 42 | >55 | Tachy-brady syndrome; Holter: AF, rate 36 to 153 bpm, frequent sinus pauses (up to 3.0 s); stress test: exaggerated rate response to exercise (max 220 bpm). On aspirin, verapamil. |
| III-1/F | 43/NA | No | SR (72 bpm) | 37 | 63 | Intermittent palpitations, triggered by stress. |
| III-2/F | 37/NA | No | SR (71 bpm) | 31 | 63 | Intermittent palpitations, triggered by stress. |
| III-3/M | 24/NA | No | SR (57 bpm) | 40 | 66 | |

AF, atrial fibrillation; APC, atrial premature contractions; dx, diagnosis; LAD, left atrial diameter; LAHB, left anterior hemiblock; LVEF, left ventricular ejection fraction; NA, not available; PPM, permanent pacemaker; RBBB, right bundle branch block; SR, sinus rhythm.

**Supplemental Table VIII**. *Drosophila* electrophysiological data.

| Fly line | No. | Resting $V_m$ (mV) | Maximum Amplitude (mV) | Peaks/ Burst | Event Duration (ms) |
|---|---|---|---|---|---|
| Heart Gal4/RNAi Cont. | 2 | -41.1 ± 0.2 | 59.6 ± 0.3 | 2.0 ± 0 | 219.9 ± 6.2 |
| Heart Gal4>*slo* RNAi | 2 | -48.3 ± 0.3 | 47.8 ± 0.4 | 3.4± 0.9 | 601.8 ± 52.8 |
| $Slo^4$/Df (3R) BSC 397 | 3 | -46.8 ± 1.0 | 72.3 ± 0.9 | 4.3± 0.3 | 860.6 ± 8.2 |
| Hand GS>*slo* RNAi+RU486 | 2 | -74.5 ± 1.4 | 64.4 ± 0.8 | 5.1± 2.5 | 490.6 ± 80.9 |
| Hand GS> *slo* RNAi No RU486 | 3 | -46.5 ± 0.5 | 60.3± 0.8 | 1.9± 0.5 | 288.6 ± 15.5 |
| Hand GS> RNAi Cont. +RU486 | 1 | -46.0 ± 0 | 58.3 ± 0.1 | 2.0 ± 0 | 394.8 ± 4.6 |
| $W^{1118}$ | 4 | -41.9 ± 0.5 | 49.6 ± 0.6 | 1.9± 0.2 | 335.0 ± 21.1 |
| Heart Gal4> $KCNMA^{wt}$;slo4/Df (3R) BSC 397 | 4 | -43.9 ± 1.2 | 55.3 ± 0.8 | 4.9± 0.5 | 902.0 ± 47.4 |
| Heart Gal4> $KCNMA^{mut}$;$Slo^4$/df | 2 | -50.4 ± 0.6 | 65.3 ± 0.5 | 5.7± 0.3 | 1033.1 ± 102.0 |

#### C. SUPPLEMENTAL FIGURE LEGENDS

**Supplemental Figure I.** Schematic of the chromosome 10q22 locus. The *KCNMA* gene is located within overlapping linkage intervals mapped in families with various cardiac and neurological disorders.

**Supplemental Figure II.** Comparison of anti-$K_{Ca}1.1$ antibodies. Immunostaining of human right ventricular tissue sections was performed using three antibodies (Alomone Labs) that target different regions of the $K_{Ca}1.1$ protein: APC-021, that binds to an intracellular epitope, residues 1097-1196; APC-151, that binds to an extracellular epitope, residues 199-213; and APC107, that binds to an intracellular epitope, residues 1184-1200. Antibodies were used alone (left column), or double-labelled with ryanodine receptor (RyR2, middle column) or connexin-43 (Cx43, right column). Images were taken on Zeiss 7DUO Confocal Microscope at magnification x20. Scale bar = 20μm.

**Supplemental Figure III.** Optimization of anti-$K_{Ca}1.1$ APC-021 antibody. Immunostaining of human right ventricular tissue sections was performed using varying concentrations of anti-$K_{Ca}1.1$ APC-021 antibody. The immunolabelling procedure was performed with omission of the primary antibody in a negative control experiment (lower right panel). Scale bar = 20μm.

**Supplemental Figure IV.** Negative controls for anti-$K_{Ca}1.1$ APC-021 antibody immunogold electron microscopy. This procedure was performed with omission of the primary antibody. In comparison to $K_{Ca}1.1$ localization demonstrated in human right atrial appendage tissue in the presence of the primary antibody (Figure 4), immunogold particles were not observed in this negative control study in (**A**) cardiomyocyte sarcomeres (T-tubules indicated by white arrow and sarcoplasmic reticulum by black arrows), (**B**) intercalated discs, (**C**) mitochondria, (**D**) coronary vascular endothelium (arrows) and (**E**) cardiac fibroblasts.

**Supplemental Figure V.** Negative control for Western blot analysis with APC-021 antibody. Nitrocellulose membrane with protein samples was blotted with a mixture of anti-$K_{Ca}$1.1 APC-021 antibody and its corresponding blocking peptide. In comparison to the $K_{Ca}$1.1 bands of 100 and 55 kDa seen with Western blotting in the presence of the primary antibody (Figure 5), addition of blocking peptide prevented antibody binding and no bands were detected. These findings confirm the specificity of APC-021 antibody binding to the $K_{Ca}$1.1 epitope.

**Supplemental Figure VI.** Restriction enzyme digestion of *KCNMA1* variation present in Family FF. The p.11-12delSSinsG sequence variation resulted in loss of an MspA1I enzyme restriction site and was evidenced by double bands on the gel. The *KCNMA1* variation was confirmed in the proband (FF-II-3, lane 2) and was also present in his father (FF-I-1, lane 3) but was not detected in his mother (FF-I-2, lane 4) or a control DNA sample (lane 5). DNA size markers were run in lanes 1 and 6; bp, base pairs.

**Supplemental Figure VII.** $K_{Ca}$1.1 in the ventricle. (**A**) Immunofluorescence analysis of human right ventricular sections shows $K_{Ca}$1.1 expression in cardiomyocytes where it colocalizes with the ryanodine receptor (RyR2) (left and center panels), and also in fibroblasts (right panel); scale bar = 5 µm. (**B**) In isolated ventricular cardiomyocytes from adult wild-type mice, $K_{Ca}$1.1 colocalizes with the ryanodine receptor-2 (RyR2; left column). $K_{Ca}$1.1 and caveolin-3 (Cav-3; right column) show co-localization intracellularly but not at the surface membrane. Merged staining was coupled with DAPI (blue) for nuclear localization; scale bar = 2.5 µm.

**Supplemental Figure VIII.** Regional differences in atrial expression of *KCNMA1* mRNA. Levels of *KCNMA1* transcript were assessed in human right atrial tissue (n=4 hearts) using qPCR. A gradient of expression was observed with relatively higher levels in the sinus node (SN) than in the paranodal (PN) region and atrial myocardium (AM). mRNA data are shown relative to 28s levels.

**Supplemental Figure IX.** Effects of age and atrial fibrillation (AF) on levels of expression of total *KCNMA1* and selected isoforms in human right atrial tissue. qPCR was performed in right atrial tissue samples from patients in sinus rhythm without a history of AF (red) and patients with documented AF (blue) aged (**A**) <70 years and (**B**) >70 years at the time of study.

**Supplemental Figure X.** $K_{Ca}1.1$ protein expression in zebrafish hearts. (**A**) Western blot analysis of $K_{Ca}1.1$ protein expression in 3 dpf zebrafish embryos injected with either control-morpholino (MO), *kcnma1a* (1a)- or *kcnma1b* (1b)-morpholino, or a combination of both (1a+1b), using an anti-$K_{Ca}1.1$ antibody that recognises both zebrafish $K_{Ca}1.1$ isoforms. Two lower bands of approximately 50 kDa were seen in the control-MO lanes but not with the *kcnma1b*-MO or when the antibody-specific blocking peptide was applied (B) and most likely represent degradation products of the *kcnma1b* isoform. (**B**) Antibody binding specificity of the non-selective anti-$K_{Ca}1.1$ antibody used in the above immunoblot was confirmed by antibody pre-incubation with antigen-specific peptide, which prevented antibody binding to both zebrafish $K_{Ca}1.1$ protein isoforms.

**Supplemental Figure XI.** Effects of *kcnma1b* knockdown on ventricular function in zebrafish. Mean data for ventricular end-diastolic diameter (EDD) and fractional shortening (FS) in control morpholino (MO)-injected and *kcnma1b* morpholino-injected embryos at 3 dpf (n=17-20 each group). Data are shown as mean ± SEM; ****$P$<0.0001 compared to control embryos.

**Supplemental Figure XII.** Cardiac abnormalities associated with *slo* genomic mutants. Flies with cardiac-specific knockdown (**A-D**) show increases in systolic interval, diastolic interval, heart period, and arrhythmia index at 3 weeks when compared to control flies. The increases in systolic and diastolic intervals and heart period were maintained at 5 weeks, however differences in the arrhythmia index were no longer seen. It is notable that control flies showed an increase in the arrhythmia index by 5 weeks of age, and mutant flies had a marked reduction in survival over this time period (**E**). We started with 200 female flies of each genotype spread over 25 flies per vial and counted the number of surviving flies after every two days.  Using a log-rank (Mantel-Cox) test, there was a significant difference in the survival curves between groups ($P$<0.0001). Flies with other mutant *slo* alleles (**F-I**) also show increases in systolic interval, diastolic interval, heart period, and arrhythmia index at 3 weeks but not at 5 weeks. (**J**) qPCR showed marked downregulation of *slo* expression in $slo^4$/Df(3R) BSC397 hearts. For all genotypes, n=14-40 each group. Data were obtained using the BioMark™ real-time PCR system (Fluidigm) and are expressed relative to wild-type (Cantonese) hearts.

**Supplemental Figure XIII.** Genomic rescue of *slo* mutants and effects of human *KCNMA1* variants. Flies with one or two copies of the genomic *slo* locus (PBac Slo/+) had increased systolic interval (**A**), diastolic interval (**B**) and heart period (**C**) at 3 weeks when compared to wild-type ($W^{1118}$) flies. The additional copies of this locus did not alter the arrhythmia index. Heart period and arrhythmia index in flies with transgenomic combinations of $slo^4$/DF(3R) BScS397, $Slo^4$ and PBac Slo/+ were similar to controls. Heart period (**E**) and arrhythmia index (**F**) were similar at 1 week and 3 weeks of age in flies with a pan-neural RNAi-mediated *slo* knockdown (Neural Gal4>slo RNAi) and in age-matched controls (Neural Gal4/RNAi control). For all genotypes, n=20-30 each group.

**Supplemental Figure XIV.** Increased arrhythmias in adult flies with cardiac-specific *slo* knockdown. Cardiac-specific *slo* knockdown in adult flies was achieved using the Hand GeneSwitch system (HandGS-Gal4). Administration of mifepristone (RU486, 100ug/mL, dissolved in ethanol) induces the HandGS-Gal4 driver, allowing temporal control of cardiac-specific expression of *slo* RNAi. **(A)** Bar graph showing percentage of control and *slo*-deficient flies that had prolonged fibrillatory events (defined by incomplete and irregular periodicity of contraction in 3 or more consecutive beats). (**B**) Mean number of prolonged fibrillatory contractions in control and slo-deficient flies. (**C**) Representative M-mode traces showing myocardial contractions in a control fly and prolonged fibrillatory activity in an adult fly with cardiac-specific *slo* knockdown (Hand GS> Slo RNAi +RU486). For all genotypes, n=30-50 each group. **$P<0.01$.

Supplemental Figure I

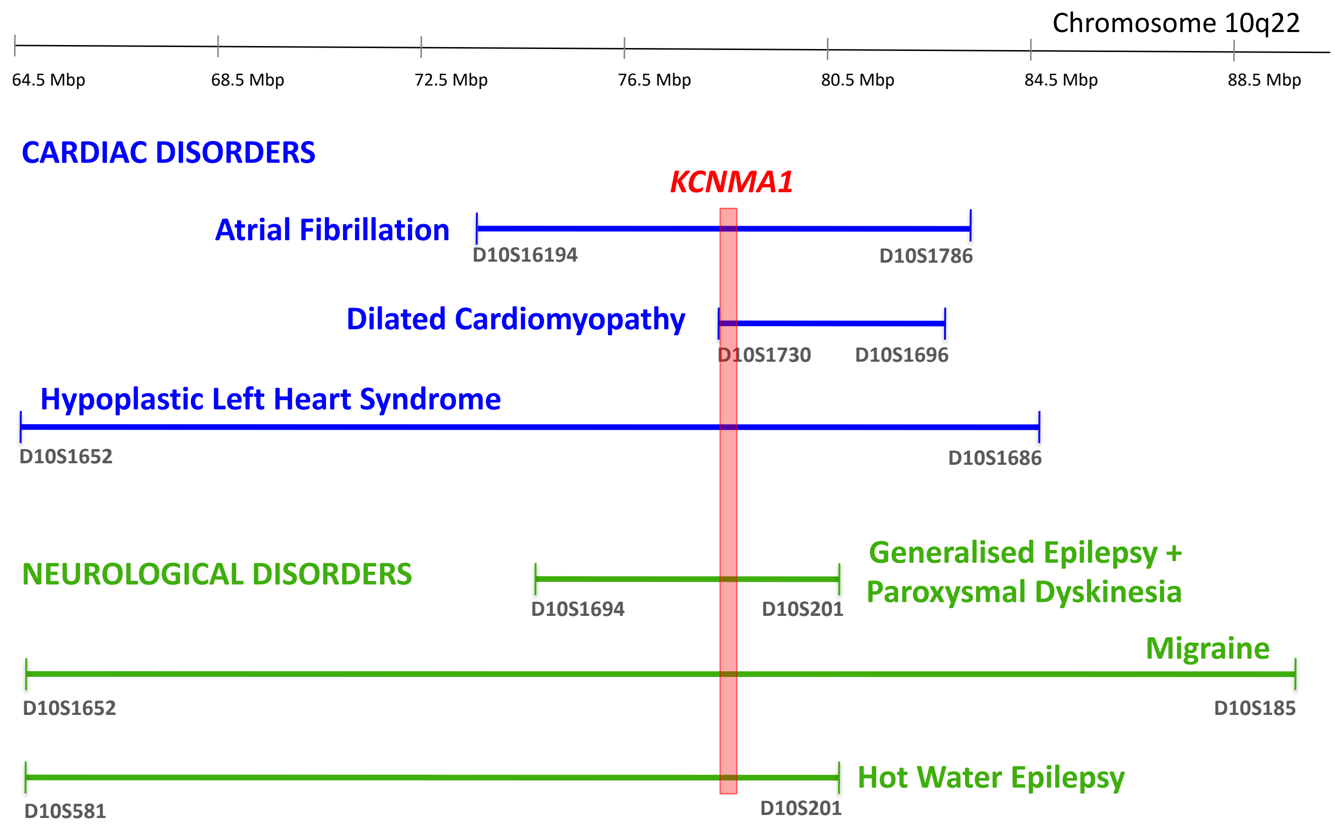

Supplemental Figure II

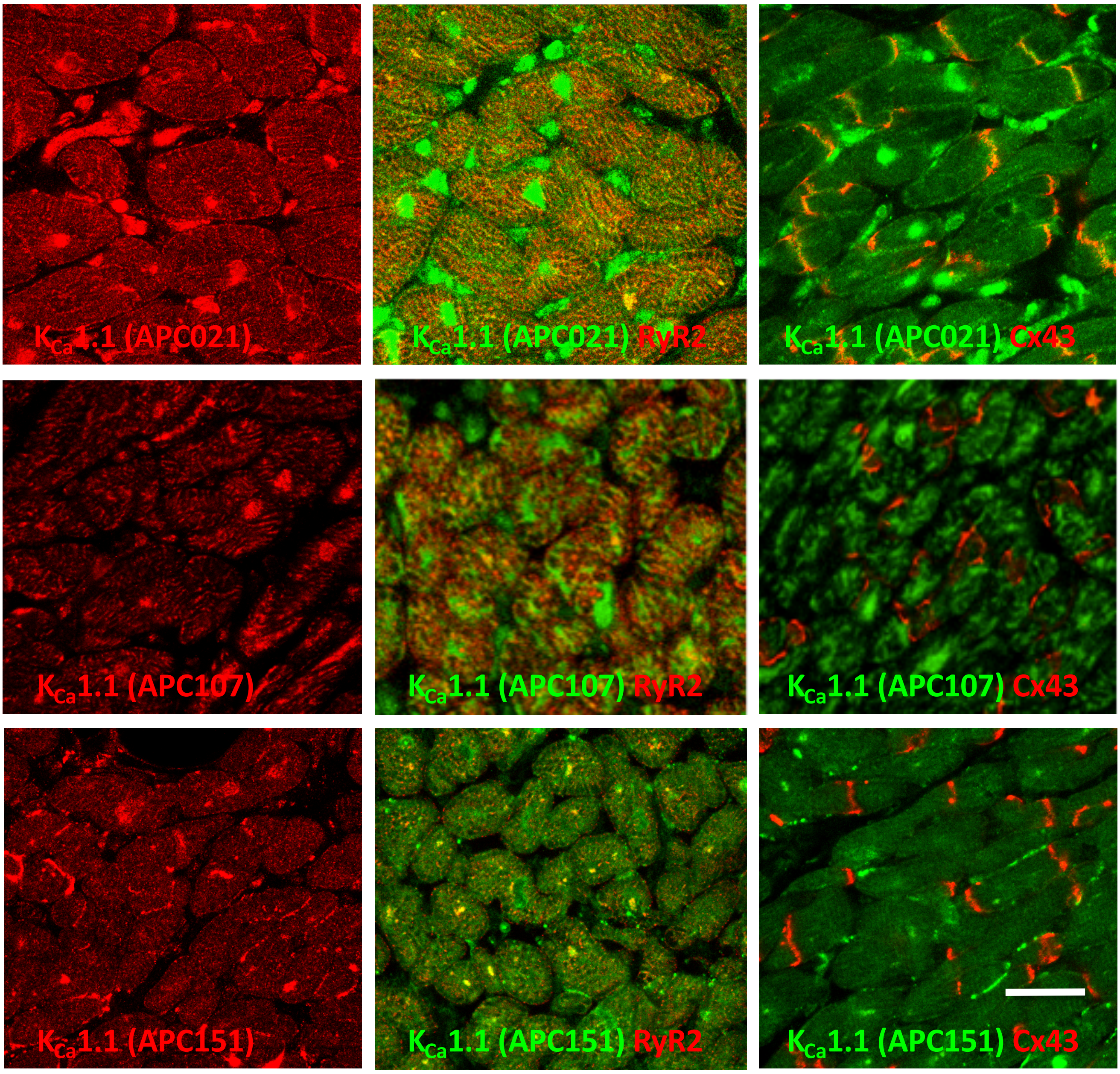

Supplemental Figure III

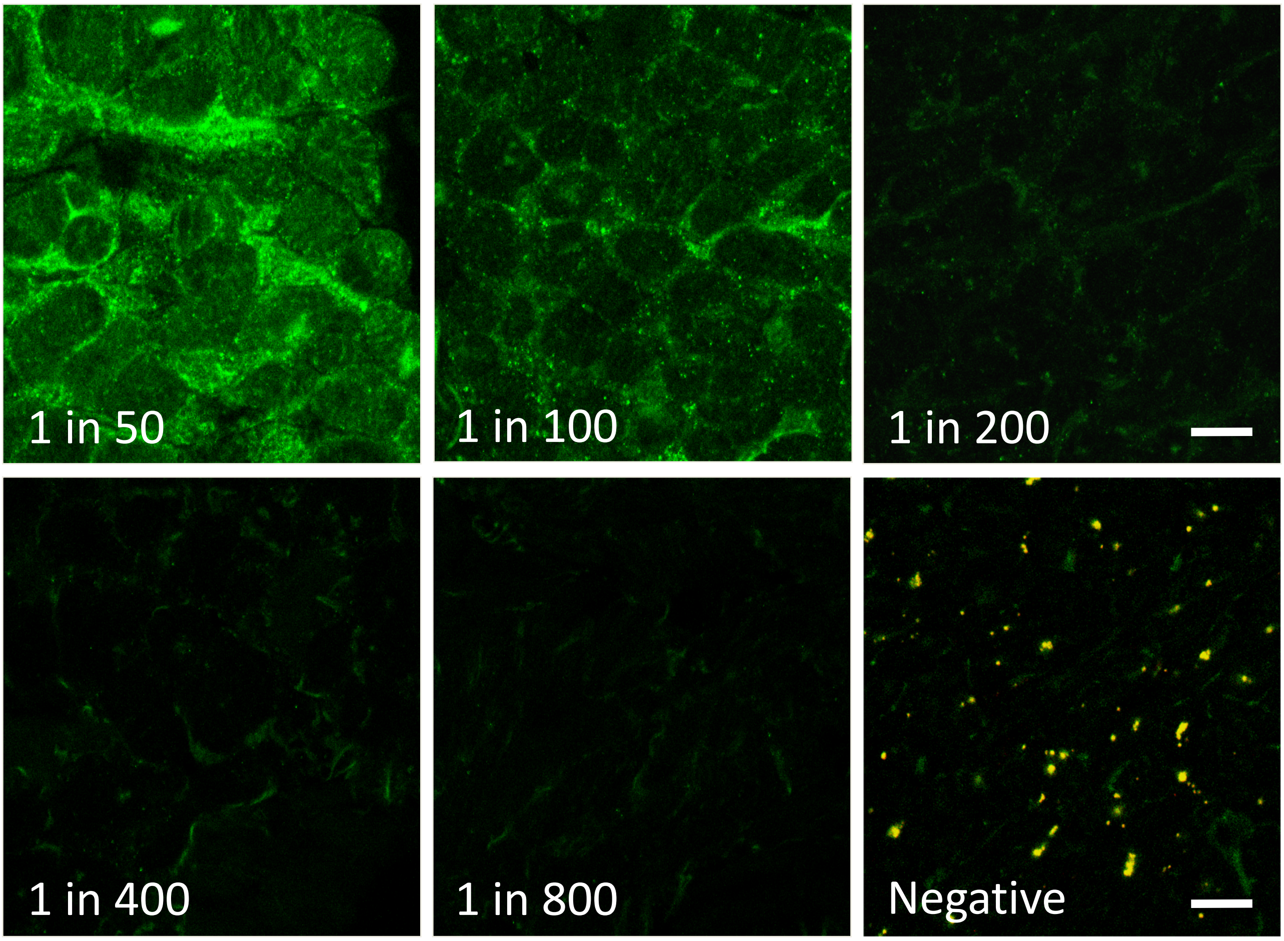

Supplemental Figure IV

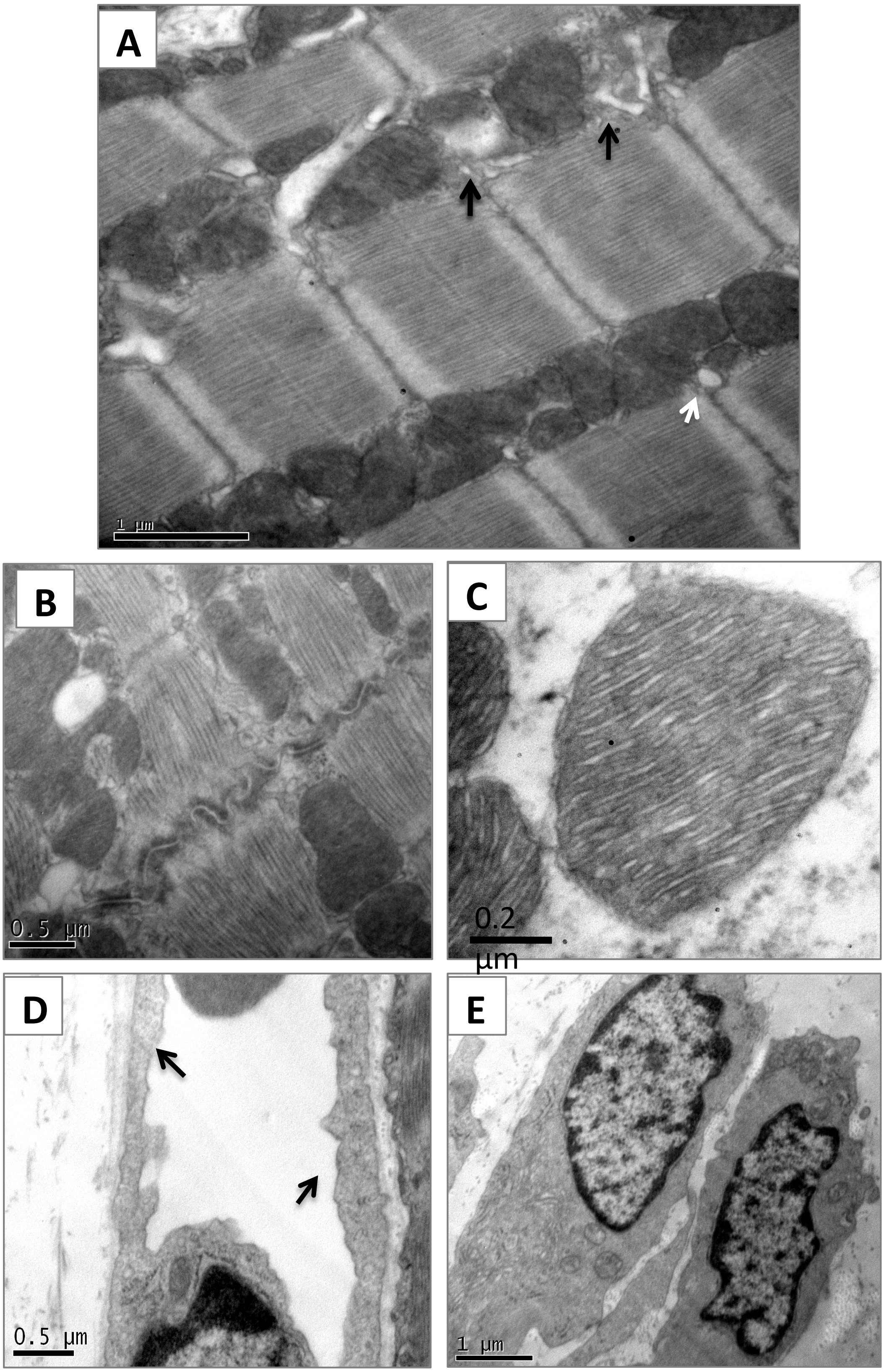

Supplemental Figure V

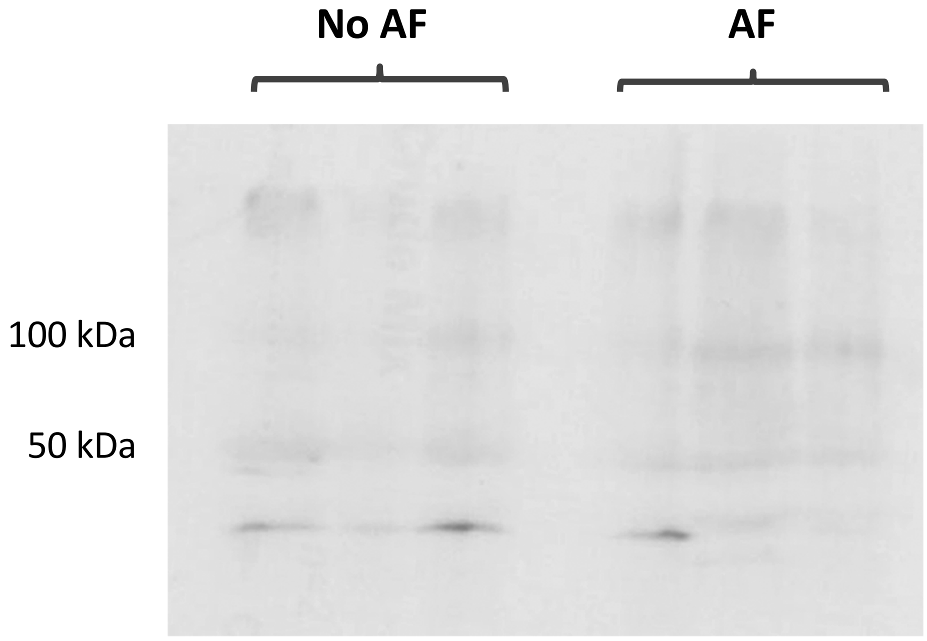

**IB: $K_{Ca}$1.1 blocking peptide + anti-$K_{Ca}$1.1 antibody**

Supplemental Figure VI

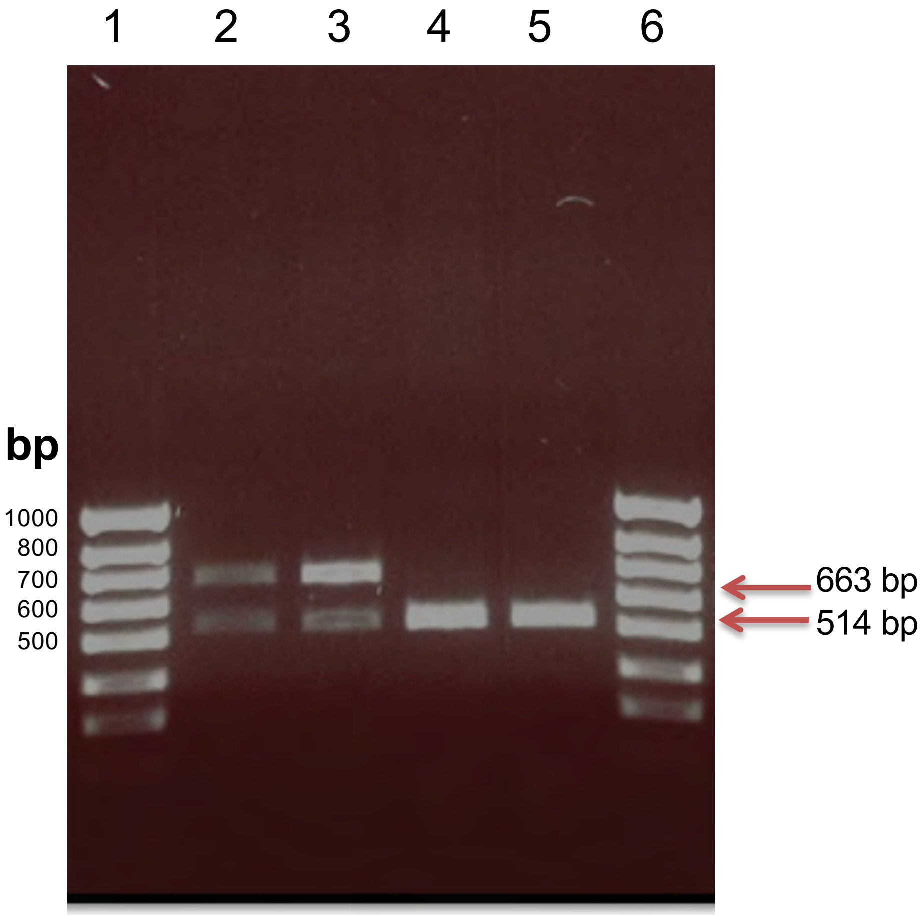

Supplemental Figure VII

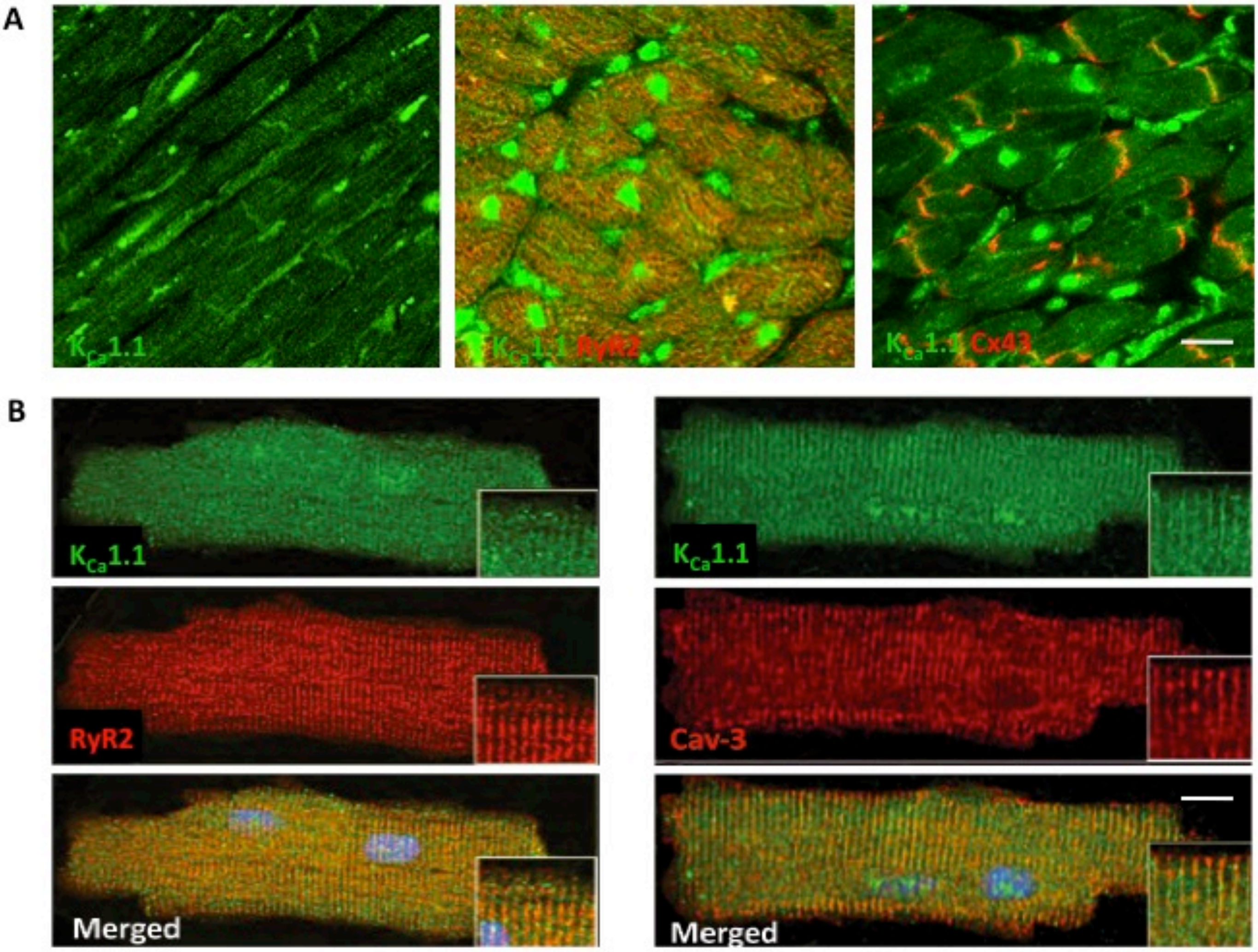

Supplemental Figure VIII

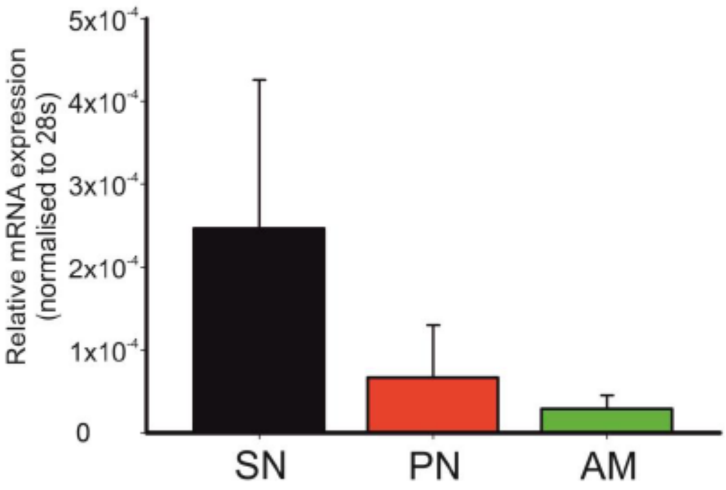

### Supplemental Figure IX

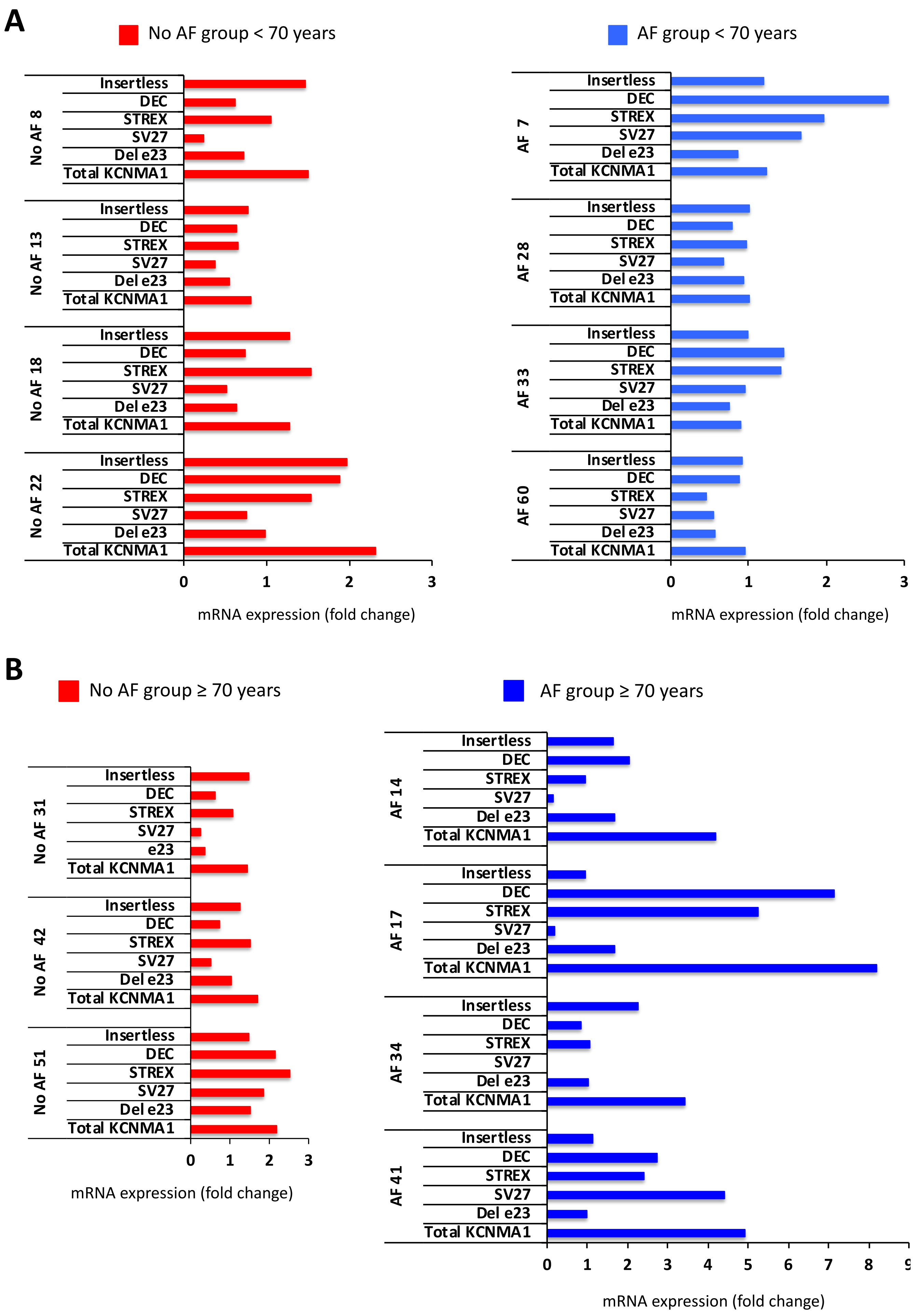

Supplemental Figure X

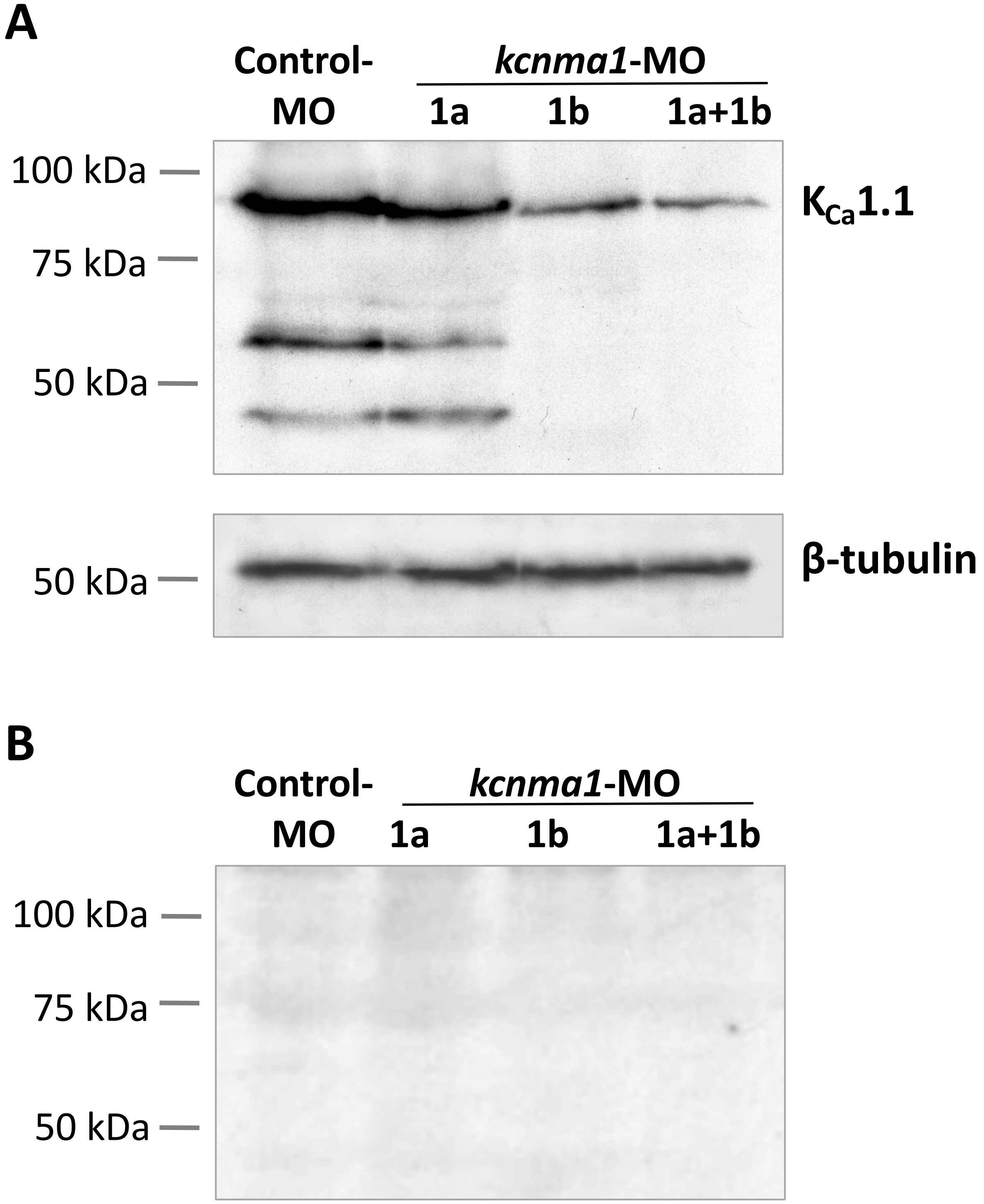

Supplemental Figure XI

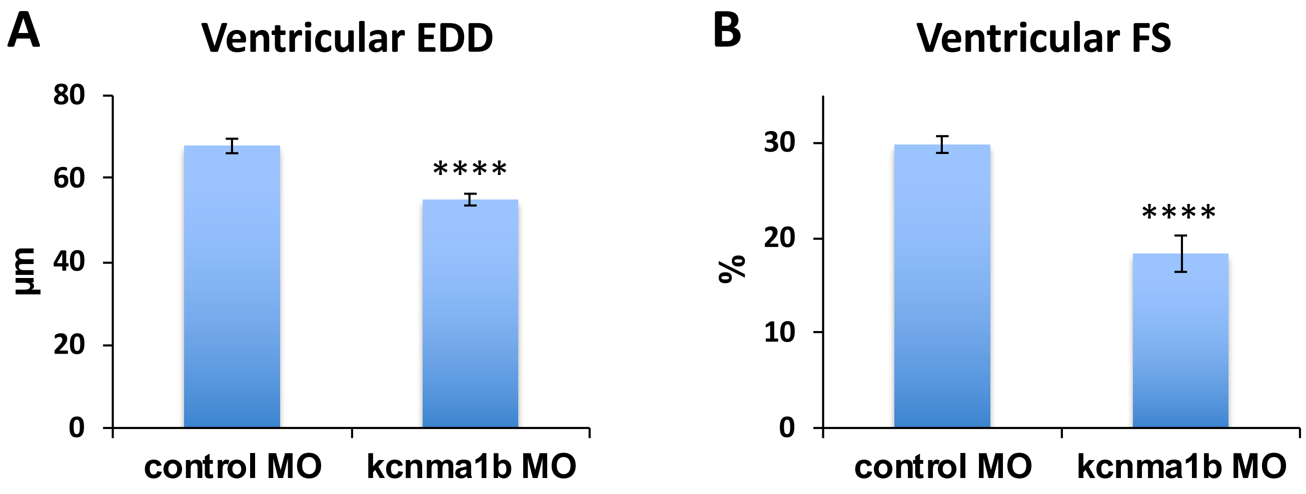

Supplemental Figure XII

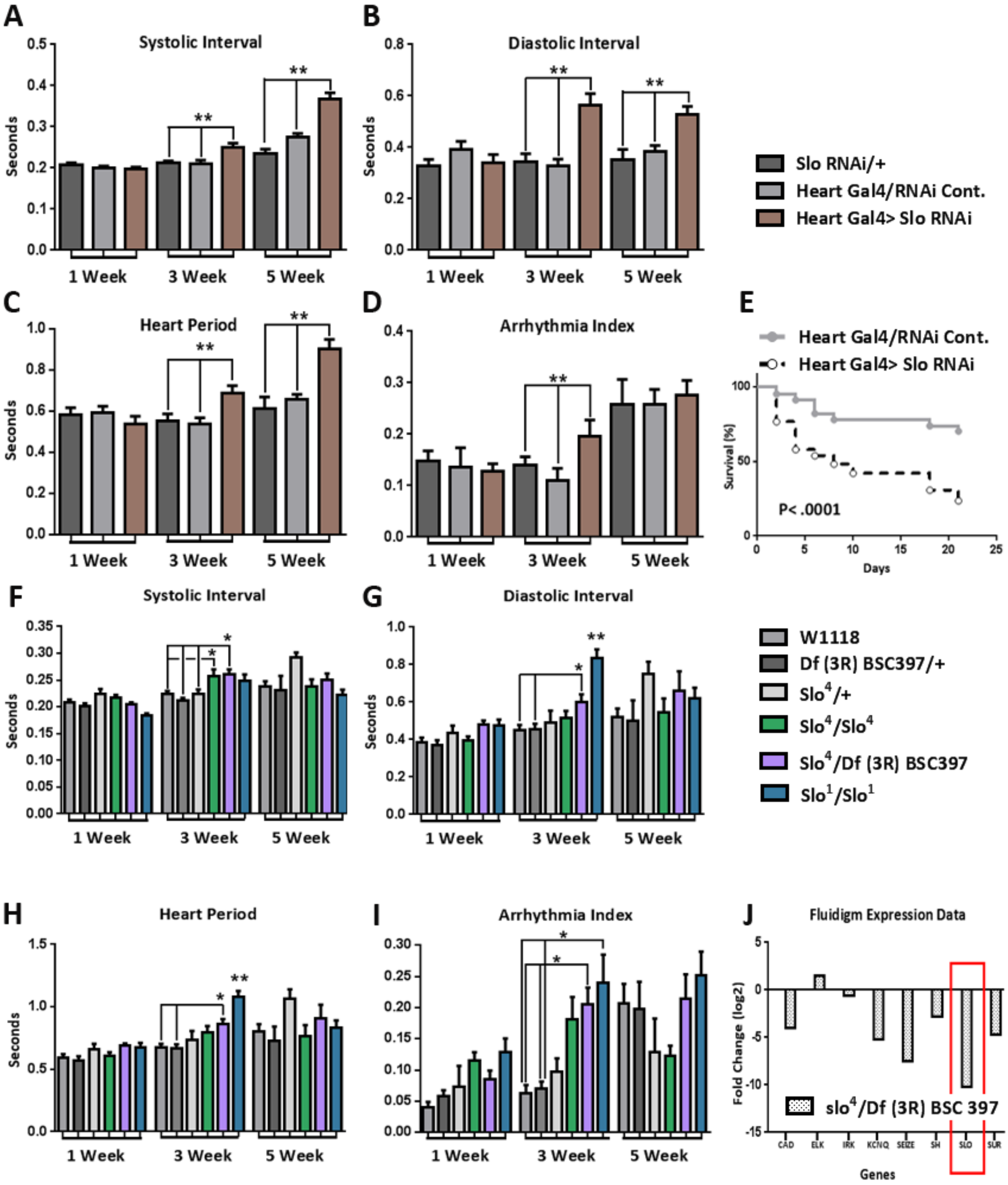

Supplemental Figure XIII

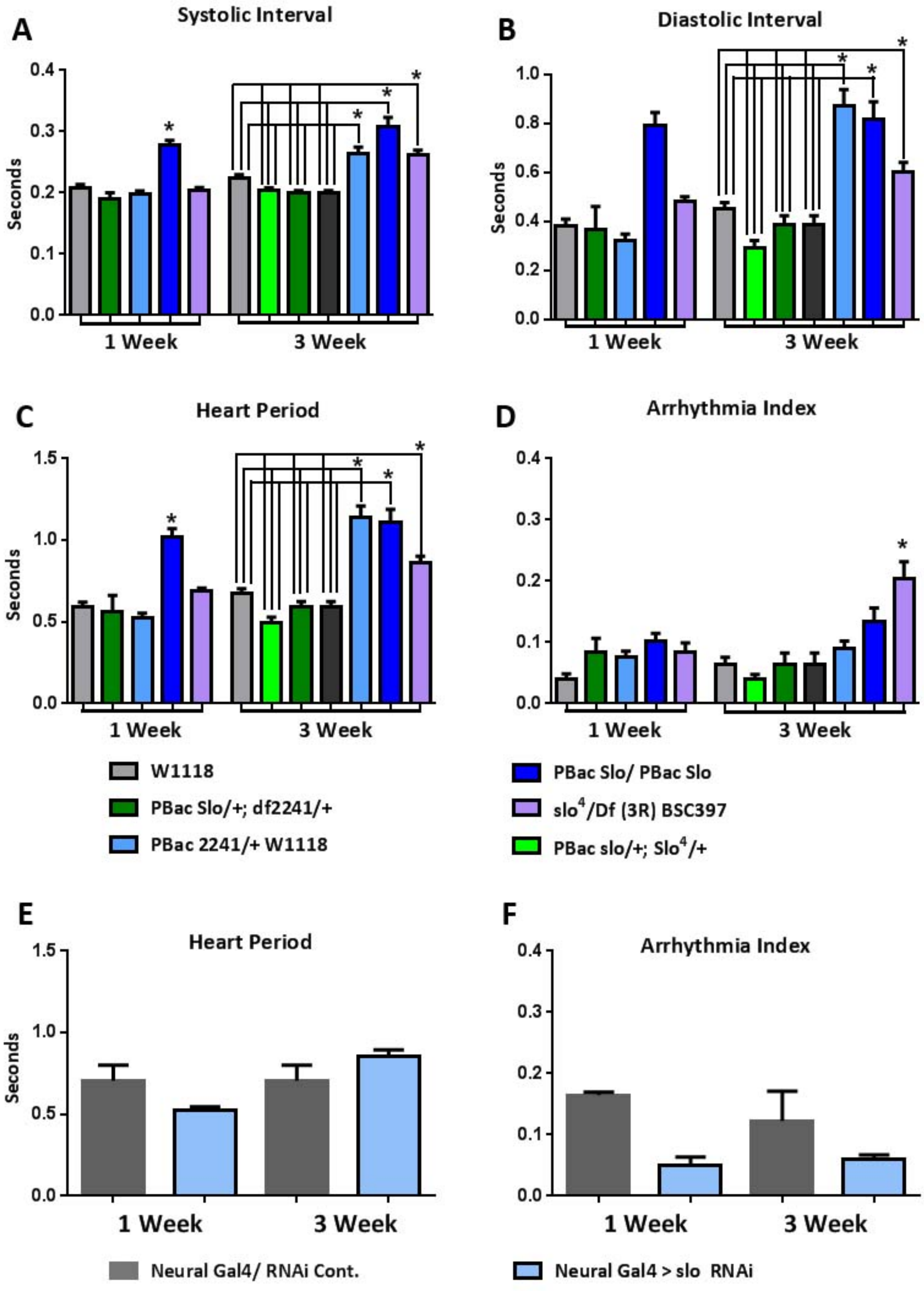

Supplemental Figure XIV

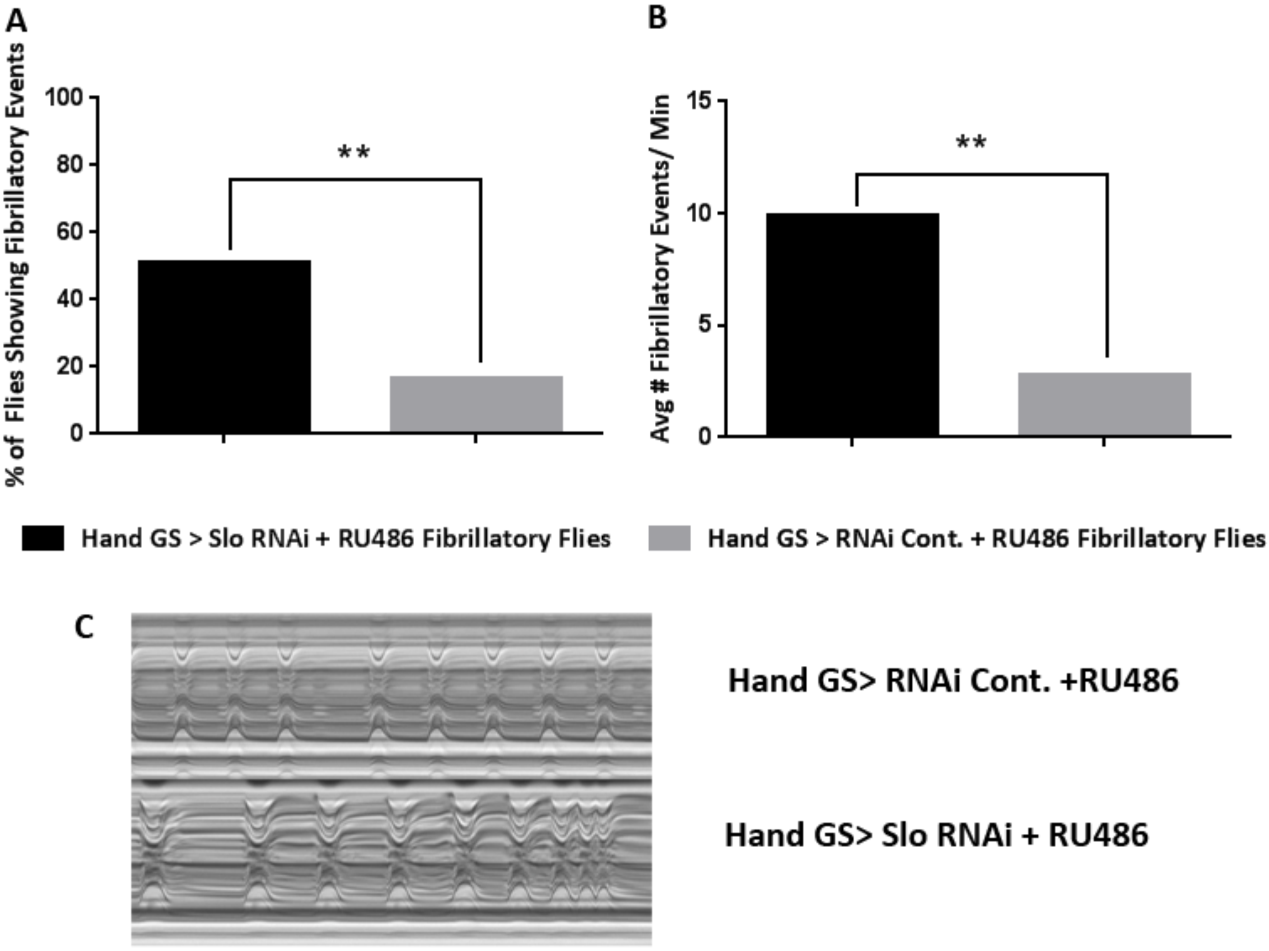

#### E. SUPPLEMENTAL REFERENCES

1. Chandler NJ, Greener ID, Tellez JO, Inada S, Musa H, Molenaar P, et al. Molecular architecture of the human sinus node: insights into the function of the cardiac pacemaker. *Circulation* 2009;119:1562-1575.
2. Lockard VG. A simple technique for post-embedding ultrastructural immunogold labelling. *Ultrastruct Pathol* 1992;16:47-50.
3. Huttner IG, Trivedi G, Jacoby A, Mann SA, Vandenberg JI, Fatkin D. A transgenic zebrafish model of a human cardiac sodium channel mutation exhibits bradycardia, conduction-system abnormalities and early death. *J Mol Cell Cardiol* 2013;61:123-132.
4. Fink M, Callol-Massot C, Chu A, Ruiz-Lozano P, Izpisua Belmonte JC, Giles W, et al. A new method for detection and quantification of heartbeat parameters in Drosophila, zebrafish and embryonic mouse hearts. *Biotechniques* 2009;46:101-113.
5. Han Z, Yi P, Li X, Olson EN. Hand, an evolutionarily conserved bHLH transcription factor required for Drosophila cardiogenesis and hematopoiesis. *Development* 2006;133:1175-1182.
6. Venken KJ, Carlson JW, Schulze KL, Pan H, He Y, Spokony R, et al. Versatile P(acman) BAC libraries for transgenesis studies in *Drosophila melanogaster*. *Nat Methods* 2009;6:431-434.
7. Osterwalder T, Yoon KS, White BH, Keshishian H. A conditional tissue-specific transgene expression system using inducible GAL4. *Proc Natl Acad Sci USA* 2001;98:12596-12601.
8. Monnier V, Iche-Torres M, Rera M, Contremoulins V, Guichard C, Lalevee N, et al. dJun and Vri/dNFIL3 are major regulators of cardiac aging in Drosophila. *PLoS Genetics* 2012;8:e1003081.

9. Ocorr K, Reeves NL, Wessells RJ, Fink M, Chen HS, Akasaka T, et al. KCNQ potassium channel mutations cause cardiac arrhythmias in Drosophila that mimic the effects of aging. *Proc Natl Acad Sci USA* 2007;104:3943-3948.
